# Serial co-expression analysis of host factors from SARS-CoV viruses highly converges with former high-throughput screenings and proposes key regulators and co-option of cellular pathways

**DOI:** 10.1101/2020.07.28.225078

**Authors:** Antonio J. Pérez-Pulido, Gualberto Asencio-Cortés, Ana M. Brokate-Llanos, Gloria Brea-Calvo, Rosario Rodríguez-Griñolo, Andrés Garzón, Manuel J. Muñoz

## Abstract

The current genomics era is bringing an unprecedented growth in the amount of gene expression data, only comparable to the exponential growth of sequences in databases during the last decades. This data now allows the design of secondary analyses that take advantage of this information to create new knowledge through specific computational approaches. One of these feasible analyses is the evaluation of the expression level for a gene through a series of different conditions or cell types. Based on this idea, we have developed ASACO, Automatic and Serial Analysis of CO-expression, which performs expression profiles for a given gene along hundreds of normalized and heterogeneous transcriptomics experiments and discover other genes that show either a similar or an inverse behavior. It might help to discover co-regulated genes, and even common transcriptional regulators in any biological model, including human diseases or microbial infections. The present SARS-CoV-2 pandemic is an opportunity to test this novel approach due to the wealth of data that is being generated, which could be used for validating results. In addition, new cell mechanisms identified could become new therapeutic targets. Thus, we have identified 35 host factors in the literature putatively involved in the infectious cycle of SARS-CoV and/or SARS-CoV-2 and searched for genes tightly co-expressed with them. We have found around 1900 co-expressed genes whose assigned functions are strongly related to viral cycles. Moreover, this set of genes heavily overlap with those identified by former laboratory high-throughput screenings (with p-value near 0). Some of these genes aim to cellular structures such as the stress granules, which could be essential for the virus replication and thereby could constitute potential targets in the current fight against the virus. Additionally, our results reveal a series of common transcription regulators, involved in immune and inflammatory responses, that might be key virus targets to induce the coordinated expression of SARS-CoV-2 host factors. All of this proves that ASACO can discover gene co-regulation networks with potential for proposing new genes, pathways and regulators participating in particular biological systems.

**Highlights:**

- ASACO identifies regulatory associations of genes using public transcriptomics data.
- ASACO highlights new cell functions likely involved in the infection of coronavirus.
- Comparison with high-throughput screenings validates candidates proposed by ASACO.
- Genes co-expressed with host’s genes used by SARS-CoV-2 are related to stress granules.

## Introduction

Genes are team players that rarely act in solitude but require the cooperation of others to carry out their physiological functions. Groups of genes are usually expressed together when necessary, under the conduction of regulatory proteins, conforming the so-called regulatory networks. In higher organisms, augmented complexity requires an increase in the number of biological functions. This is mainly achieved by expanding regulatory relationships rather than the number of participating genes (Long et al., 2016; Rogers and Bulyk, 2018). The combinatorial or regulatory activity of proteins on structural genes seem to account for the genetic plasticity required for development, organ functional diversity, responses to environmental changes and other emergent properties of multicellularity (Lobo, 2008; Spitz and Furlong, 2012). One gene might play different roles in different regulatory contexts, but it will always be accompanied by other genes involved in that function. We hypothesize that, even under this complex scenario, this regulatory linkage among proteins sharing biological functions and pathways can be brought to light by carefully analysing co-expression profiles from functional genomics and transcriptomic experiments.

The genomics era is generating a wealth of information about gene expression in many different biological processes and experimental approaches. Although designed to address specific questions, genomic expression data (as those from microarrays or RNA-Seq experiments), can provide a huge amount of information regarding regulatory relationships, usable to address different unrelated problems. One of the main obstacles encountered when analysing data from different experimental sources is standardization. Lately, curated databases have been deployed to gather and normalize genomics experimental data. Expression Atlas is one of them (Papatheodorou et al., 2020). As of July 2020, it holds normalized data from 1403 human transcriptomics experiments. These data contain valuable information on regulatory relationships that can be exploited to gain insights into any human biological system.

With all this in mind, we have devised a bioinformatics method based on an algorithm called Automatic and Serial Analysis of CO-expression (ASACO) that analyses information of multiple human transcriptomics experiments from Expression Atlas to predict novel regulators and functional partners of a given function. To challenge this algorithm, we have addressed the analysis of several known human proteins that are important to Severe Acute Respiratory Syndrome Coronavirus 2 (SARS-CoV-2) infection cycle and predicted new possible partners and regulators of these functions. The rationale behind this choice is the amount of experimental high-throughput genomic data, which is being produced because of the interest generated by the COVID-19 pandemics, providing an excellent opportunity to benchmark our *in silico* output. Moreover, we must also value the possibility of generating new knowledge on SARS-CoV-2 infection process. The virus infection cycle rests on the host’s physiological functions to take place, requiring the sequestration or co-option of multiple host factors. In the first place, it is clearly stated the general inhibition activity exerted by SARS viruses on host’s mRNAs translation (Kamitani et al., 2009; Thoms et al., 2020). But also other aspects of the host physiology are used or modified by the virus, such as the general RNA metabolism, protein synthesis, cell cycle regulation, cell membrane dynamics, autophagy, apoptosis, unfolded protein response, MAP kinases, innate immunity and inflammatory response (Fung and Liu, 2018; Sanche et al., 2020). The infection cycle heavily relies on tight interactions with the host endomembrane organelles. Coronaviruses enter the host cell by an endocytic mechanism involving the fusion between its coat membrane and the cell plasma membrane. This fusion is mediated by plasma membrane receptor recognition through the virus spike (S) protein (Belouzard et al., 2012; Boulant et al., 2015; Heald-Sargent and Gallagher, 2012). Moreover, interaction with endoplasmic reticulum-derived membrane structures is essential for gene expression and proper nucleocapsid formation. Both posttranslational modifications (Fung and Liu, 2018) and folding assistance by cell chaperones are important for correct virus functions (Fukushi et al., 2012). Virus assembly takes place on the endoplasmic reticulum▫Golgi intermediate compartment (ERGIC), where they are engulfed in membrane-bound vesicles to be finally secreted out of the host cell (Fung and Liu, 2019; Masters, 2006).

Understanding the basis of interactions of these pathways is crucial to fight infection. Translatomics and proteomics analysis of the response to viral infection have recently revealed cellular functions possibly involved in the infection process (Bojkova et al., 2020). In the same way, interactomics using viral proteins as baits to find human proteins physically interacting with them, have also revealed possible new therapeutic targets (Gordon et al., 2020). Even more complex network analysis combining protein-protein interactions and transcriptomics pointed to new potential targets as well (Yadi Zhou et al., 2020). It is also noteworthy a study made on SARS-CoV knocking down kinases by sRNAi treatment. This study highlighted cellular kinases that are important to SARS-CoV virus infection and possibly crucial for SARS-CoV-2 as well (de Wilde et al., 2015). Notably, not only the cellular but also the systemic response to the virus infection is important, and it seems to play a relevant role in SARS-CoV-2 infection as in many other viral cycles. Evidence is arising suggesting that SARS viruses are able to exploit the organism’s innate immune response on its own benefit, co-opting some components of the response, as stress granules and processing bodies (Fung et al., 2016; Moosa and Banerjee, 2020; Sola et al., 2011), and recruiting as host factors those genes co-expressed with the innate immune response (Pinto et al., 2020; Sungnak et al., 2020; Ziegler et al., 2020). This evidence stresses the necessity of research to a broader systemic landscape to explain, for instance, SARS-CoV-2 influence on the inflammatory response (Giamarellos-Bourboulis et al., 2020; Huang et al., 2020; Yonggang Zhou et al., 2020).

To help identifying functional companions in regulatory networks for a given gene we have refined our algorithm to search for genes co-expressed with that initial gene along many unrelated transcriptomic experiments. We have applied this analysis to 35 human genes experimentally shown to play a role in SARS-CoV and/or SARS-CoV-2 infections. Here, we present the results of this test, compare with those from several experimental high-throughput genomic analysis and suggest some new functions possibly involved in SARS-CoV-2 biology.

## Materials and methods

### Seed and expression data collecting

We searched in PubMed for human genes involved in the infection cycle of SARS-CoV-2 and SARS-CoV, and both the gene names and UniProtKB entry identifiers were collected. We called these genes as seeds. Then, we used the gene name in the Expression Atlas database to download the experiments where the seed was differentially expressed. This database provides 1352 standardized human transcriptomics experiments with a total of 3744 comparisons of different biological conditions coming from published results (Papatheodorou et al., 2020). Then, we used a program written in Python language to get the expression matrices from all the considered experiments. These matrices contain the logarithm in base 2 of the fold change value (log2FC) for each gene within the experiments. We took the log2FC from all the genes with this value higher than 1 or lower than -1, and p-value lower than 0.05, in at least one of the collected experiments, and created a matrix of log2FC with the genes and the experiments. Finally, the Pearson correlation coefficient was independently calculated with the expression profile, for all the experiments, between the seed and each of the other genes.

### ASACO algorithm

The ASACO algorithm (Automatic and Serial Analysis of CO-expression) involves a methodology to select genes that share similar behavior in terms of expression with the seed. The algorithm is based on fold change signs, as it is described as follows. The procedure begins with the matrix of fold change values extracted from the experiments of the Expression Atlas database in which the seed appears with a fold change value.

First, we only consider experiments where the absolute value of fold change for the seed is equal or higher than 1, in order to consider only experiments with significant expression changes. Let be *s*_*j*_ with 1≤*j* ≤ *m* the fold change of the seed for the experiment j and m is the number of experiments after the above-mentioned removal. Therefore, |*s*_*j*_| ≥1, ∀*j*: 1 ≤ *j* ≤ m.

The remaining genes which appears at least once in any of the experiments are considered to select co-expressed genes with the seed. Let be *G*_*ij*_ the matrix of fold changes of those genes, with 1 ≤ *i* ≤ *n* and 1 ≤ *j* ≤ *m*. Specifically, let be *G*_*ij*_ the fold change of the gene i in the experiment j, being n the number of genes.

The following metrics were defined for each gene *G*_*i*_. First, *P*_*i*_ is the proportion of fold changes of the gene *G*_*i*_ that have the same sign than the seed in the same experiments. *P*_*i*_ is defined as it is shown in Equation 1. It is assumed that the function sign() returns 1 if the sign of its operand is positive, and -1 otherwise.

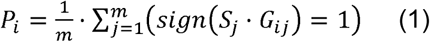

In second place, *N*_*i*_ is defined as the proportion of fold changes of the gene *G*_*i*_ that have distinct sign than the seed in the same experiments. *N*_*i*_ is defined as it is shown in Equation 2.

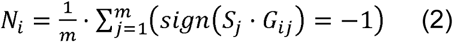

Then, *Z*_*i*_ is the proportion of experiments in which the gene *G*_*i*_ does not have any fold change annotated in the database. *Z*_*i*_ is defined as it is shown in Equation 3.

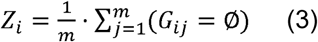

As metrics were defined, *P*_*i*_ + *N*_*i*_ + *Z*_*i*_ = 1 for 1 ≤ *i* ≤ *n*. Then, a p-value *P*(*P*_*i*_) was computed for each *P*_*i*_ from the sample distribution of *P*_*i*_. Similarly, a p-value *P*(*N*_*i*_) was computed for each *N*_*i*_ from the distribution of *N*_*i*_.

The p-value of *P*_*i*_, *P*(*P*_*i*_), is the probability that there are values greater than or equal to *P*_*i*_ in the distribution of P, given the sample obtained from the experiment data base. Analogously the p-value of *N*_*i*_, *P*(*N*_*i*_), is the probability that there are values greater than or equal to *N*_*i*_ in the distribution of *N*.

Finally, selected genes were divided into two groups: a) those that are directly correlated in terms of the sign of their fold changes, and b) those that are inversely correlated.

The first group is defined as *D* = {*G*_*k*_} for each k such that *P*(*P*_*k*_) ≤ .01 and 1 ≤ *k* ≤ |*D*|, where |*D*| is the number of selected directly correlated genes. Analogously, the second group of selected genes is defined as I = {*G*_*k*_} for each k such that *P*(*N*_*k*_) ≤ .01 and 1 ≤ *k* ≤ |*I*|, where |*I*| is the number of selected inversely correlated genes.

In this way, selected directly correlated genes are those which fold change signs are mostly the same than the seed in the same experiments, since their value of P is significantly high and, therefore, the probability of finding genes with a higher value of P is very low (≤ .01). Analogously, selected inversely correlated genes are those which fold change have mostly the opposite sign than the seed in the same experiments, since their value of N is significantly high and, therefore, the probability of finding genes with a higher value of *N* is very low.

The ASACO algorithm is written in R language and available at https://github.com/UPOBioinfo/asaco

### Functional annotation and pathway analysis

Functional annotation for both seeds and co-expressed genes were obtained from Biomart (Durinck et al., 2009). The functional enrichment were made with KEGG Pathway (Kanehisa et al., 2017), and Reactome (Jassal et al., 2020), using the R libraries biomaRt, clusterProfiler, and ReactomePA, and a p-value cutoff of 0.05.

To find pathways with a significant averaged correlation with the seeds, all human genes were grouped by the Reactome pathway where they belong. A Wilcoxon test was calculated by each pathway and the p-value was adjusted by the FDR method. Finally, only the pathways with p-value equal or lower than 1e-05, and a median correlation higher than the third quartile of the distribution of all the analyzed genes were considered as significant pathways.

To discover transcription factors into the gene datasets, the gene ontology term ‘DNA-binding transcription factor activity’, together with those of ‘regulation of DNA-binding transcription factor activity’ were searched (GO:0003700, GO:0051090, GO:0051091, GO:0043433).

### Comparison with high-throughput experiments

Supplementary files with genes from the published interactome of SARS-CoV-2 (Gordon et al., 2020) and both translatome and proteome (Bojkova et al., 2020) were downloaded, and gene identifiers were mapped to UniProt accession numbers. Then, co-expressed proteins from ASACO were independently compared to every dataset. To calculate the number of expected matches we take 21489 as the number of total proteins in the human proteome, based on the Biomart gene type ‘protein_coding’. To calculate the p-value of the number of found matches, a hypergeometric test was used (dhyper function in R).

### Genes regulated by interferon and genes related to stress granules

To check if a gene was induced or repressed by interferon, its expression was evaluated using the Interferome database v2.01 (Rusinova et al., 2013). Only experiments where a gene had a fold change higher than 2 were considered. When a gene appeared differently expressed in more than one experiment, the average value was calculated.

Genes related to stress granules were extracted from the MSGP database (Mammalian Stress Granules Proteome), that store a total of 464 proteins (Nunes et al., 2019). Available gene names were mapped to UniProt accession numbers.

## Results

### Genes involved in well-known pathways from host factors show a positive correlation to these factors over hundreds of experiments

A set of human genes are known to be involved in the infectious cycle of SARS-CoV and SARS-CoV-2. We searched the literature and found 35 protein-coding genes that participate in different stages of their infection (Table 1). Then, we classified those host factors by the viral activity where they are involved (entry, replication, or vesicle fusion), and created two subgroups according to their type of function (proviral or antiviral). Proteins encoded by these genes show common cellular functions (Fig. 1). The three most populated groups of genes encode for: five replication proteins involved in metabolism of RNA, mainly mRNA splicing (DDX1, DDX5, HNRNP1, PPIH, and ZCRB1), three proteins of vesicle fusion involved in endoplasmic reticulum to Golgi apparatus trafficking (COPB1, COPB2, and GBF1), and a large number of proteins involved in immune system, mainly interferon and cytokine signaling. This latter group includes five antiviral proteins (EIF2AK2, BST2, IFITM1, IFITM2, and IFITM3), together with the replication proteins IMPDH1, IMPDH2, and PPIA, and the vesicle protein VAPA. Furthermore, around the proteins of this group appear several proteins relevant for the virus entry into the host cell such as the proteases FURIN and cathepsin L1 (CTSL), as well as BSG and PIKFYVE. In fact, CTSL that improves the efficiency of the viral entry, is known to be involved in both adaptive and innate immune system (Zavašnik-Bergant et al., 2004).

**Table 1.**
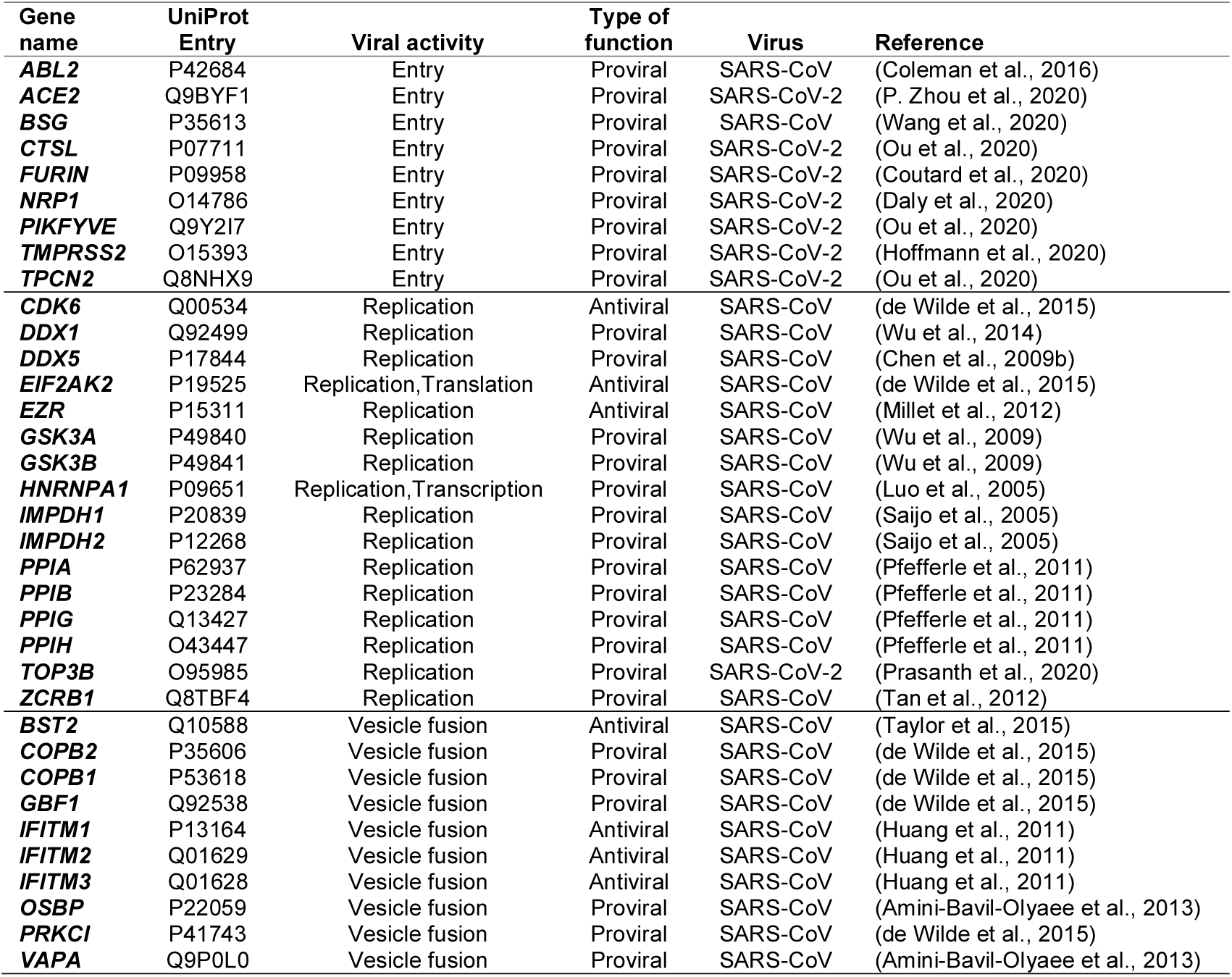
Gene targets which are expected to be essential for the infectious cycle of SARS-CoV-2. Human genes used for the ASACO analysis. Their probable activity and the type of function observed in the tested virus are indicated.

**Fig. 1.**
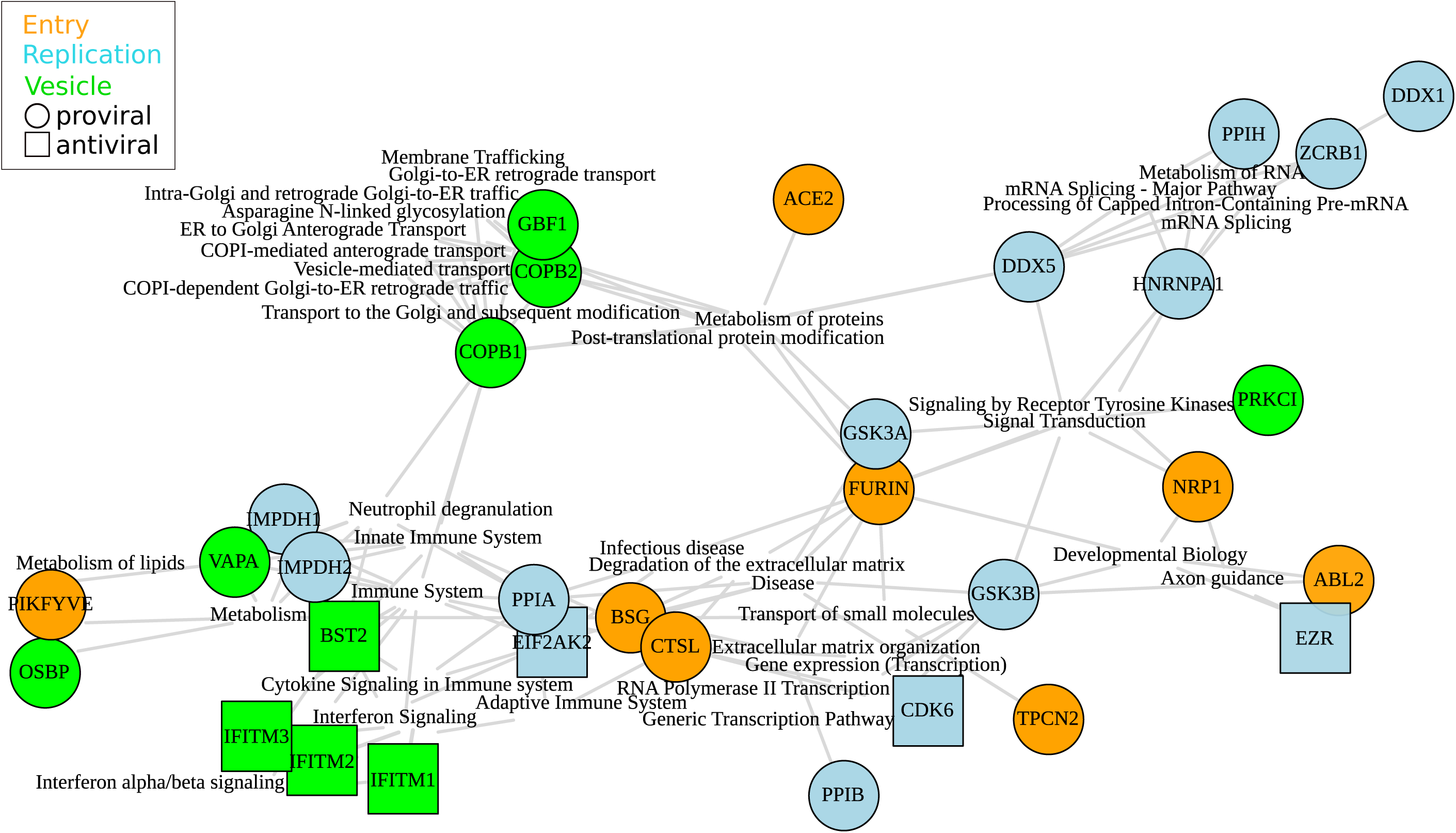
Reactome pathways shared between three or more seeds. Seeds are highlighted by its type of activity (proviral or antiviral), and its function in the infection (entry, replication, or vesicle fusion). Seeds with none shared pathways are not showed (PPIG, TOP3B, TMPRSS2).

All these proteins are part of autochthonous cell processes where they interact with others. Those proteins involved in the same biological processes frequently share expression profiles, which suggests they are co-regulated, often even showing common regulators. To study the expression relationships between the previous host factors and their co-expressed genes, we obtained transcriptomics experiments where they appear to be differentially expressed (Fig. 2). Starting from these host factors, that we named as seeds from now on, a total of 1381 different experiments were analyzed, 116 of them related to conditions involving viruses different from SARS-CoV-2. Then, the correlation value between the expression profile of these seeds versus all the other human genes through the different experiments was calculated.

**Fig. 2.**
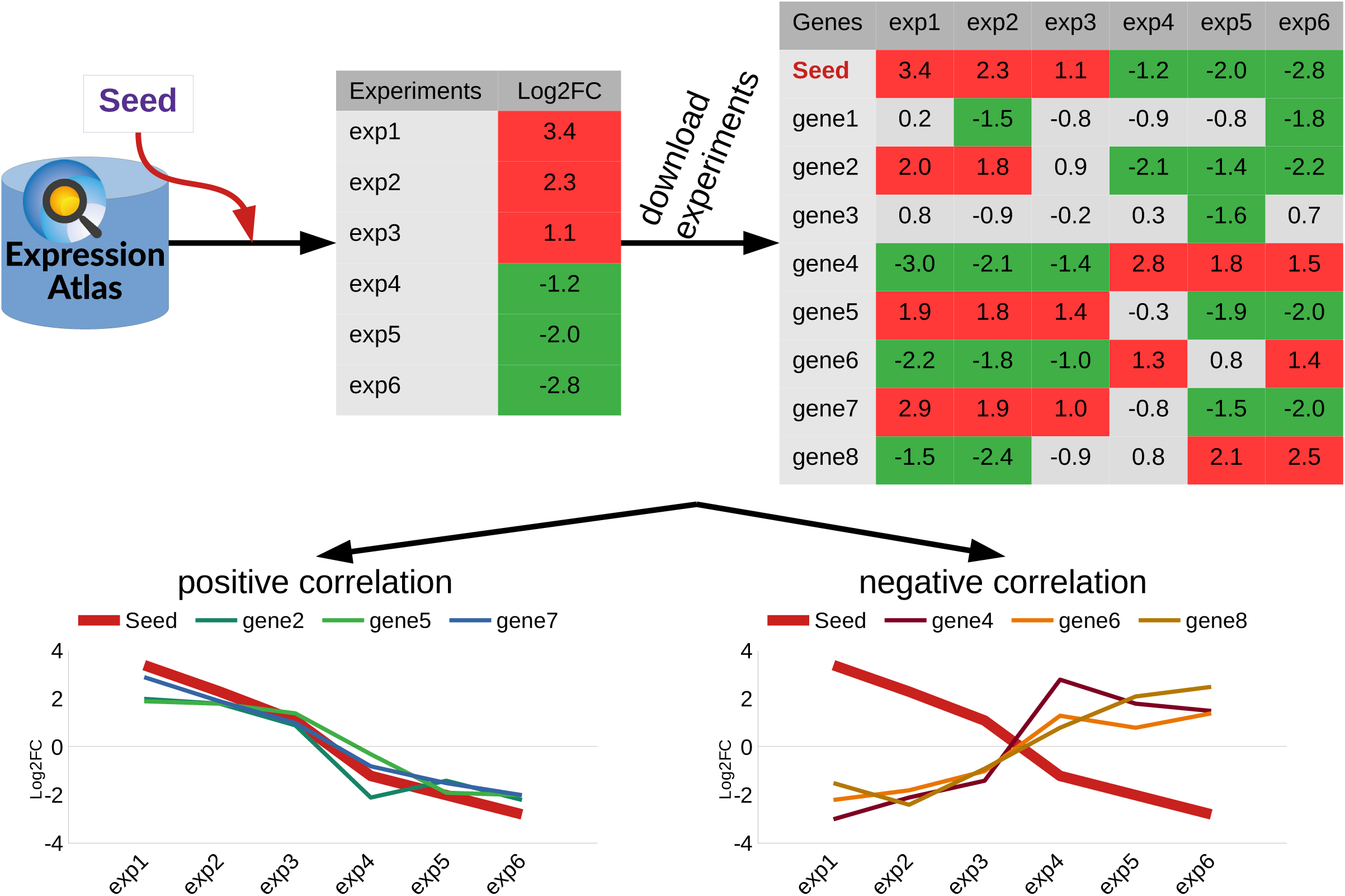
Workflow followed by ASACO; Automatic and Serial Analysis of CO-expression. Transcriptomics experiments where seeds are differently expressed are searched in the Expression Atlas database. Different experiments can be found for a given seed, along with its fold change value (log2FC). Then, experiments are downloaded and the complete expression matrix by experiment is obtained. In this matrix, other genes can be differentially expressed and their log2FC are also extracted. The log2FC values are used to create expression profiles for each gene. When the expression profiles have a positive correlation with that of the seed, their corresponding genes are expected to be co-regulated and functionally related to the seed (gene 2, gene 5, gene 7), and when the expression profiles have a negative correlation with that of the seed (gene 4, gene 6, gene 8), they are expected to have an inverse behavior in terms of expression.

It is expected that genes participating in the same biological process than the seed show a high correlation with it. Thus, we separated the correlation values for all the genes versus a given seed by the pathway they belong, and those pathways with a high average correlation value were analyzed. As a result, the most significant obtained pathways were usually those corresponding to the ones already annotated for the corresponding seed (Fig. 3a), which would support the previous assumption that genes participating in the biological process where the seed is involved show a positive correlation with the seed. In addition, the most significant pathways found when analyzing all the seeds are once again discriminating those involved in replication (mainly metabolism of RNA, and cell cycle) from those related to vesicle fusion, including the main five antiviral genes (Fig 3b). Remarkably, seeds involved in cell entry appeared linked to these groups of genes. TPCN2 and BSG are linked to the group of replication seeds by the mitochondrial translation pathway, while NRP1 and CTSL appear linked to the group of antiviral genes, once again by both interferon and cytokine signaling pathways (Fig. 3b). These two groups would highlight the two important cell interactions with the virus: functions essential for its infectious cycle such as mRNA splicing, and the cytokine antiviral response.

**Fig. 3.**
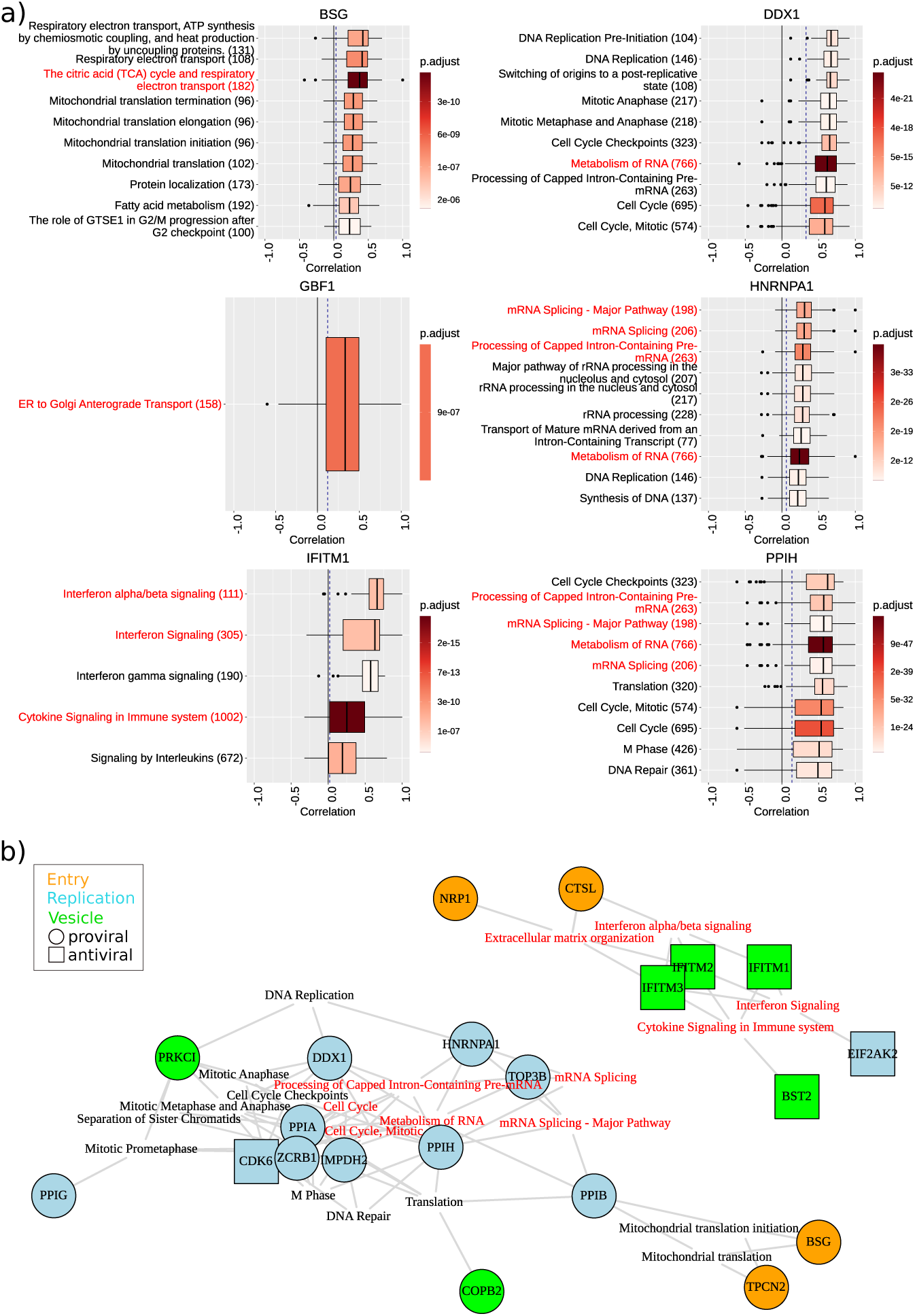
Significative pathways found by ASACO. **a)** Distribution of correlation values of genes belonging to the most significant pathways (maximum 10) for 6 of the seeds. Pathways already annotated for the seed are highlighted in red color. The number of the genes that the pathway has annotated is shown in brackets. The solid line marks correlation 0, and the dashed line marks the median of the correlation for all the genes. **b)** Significative pathways shared between three or more seeds. Seeds are highlighted by its type of activity (proviral or antiviral), and its function in the infection (entry, replication, or vesicle fusion). Seeds with none shared pathways are not showed (ABL2, ACE2, COPB1, DDX5, EZR, FURIN, GBF1, GSK3A, GSK3B, IMPDH1, OSBP, PIKFYVE, TMPRSS2, VAPA). Pathways already annotated for any seed are highlighted in red color.

### Genes with expression profiles like the seed ones present common pathways as well as others related to viral infections

To assess the agreement of functions from the best correlated genes to the ones of seeds, the expression profile of each seed was constructed using the available transcriptomic conditions. Genes with a similar expression profile (co-expressed genes) as well as genes with an inverse profile (inversely expressed genes) were obtained (Fig. 4). The total number of found co-expressed genes was 2567, although several of them were common to different seeds. Thus, the number of different co-expressed genes was 1899, while the number of the inversely expressed ones was 1578 (Suppl. Fig. S1, Suppl. Table S1).

**Fig. 4.**
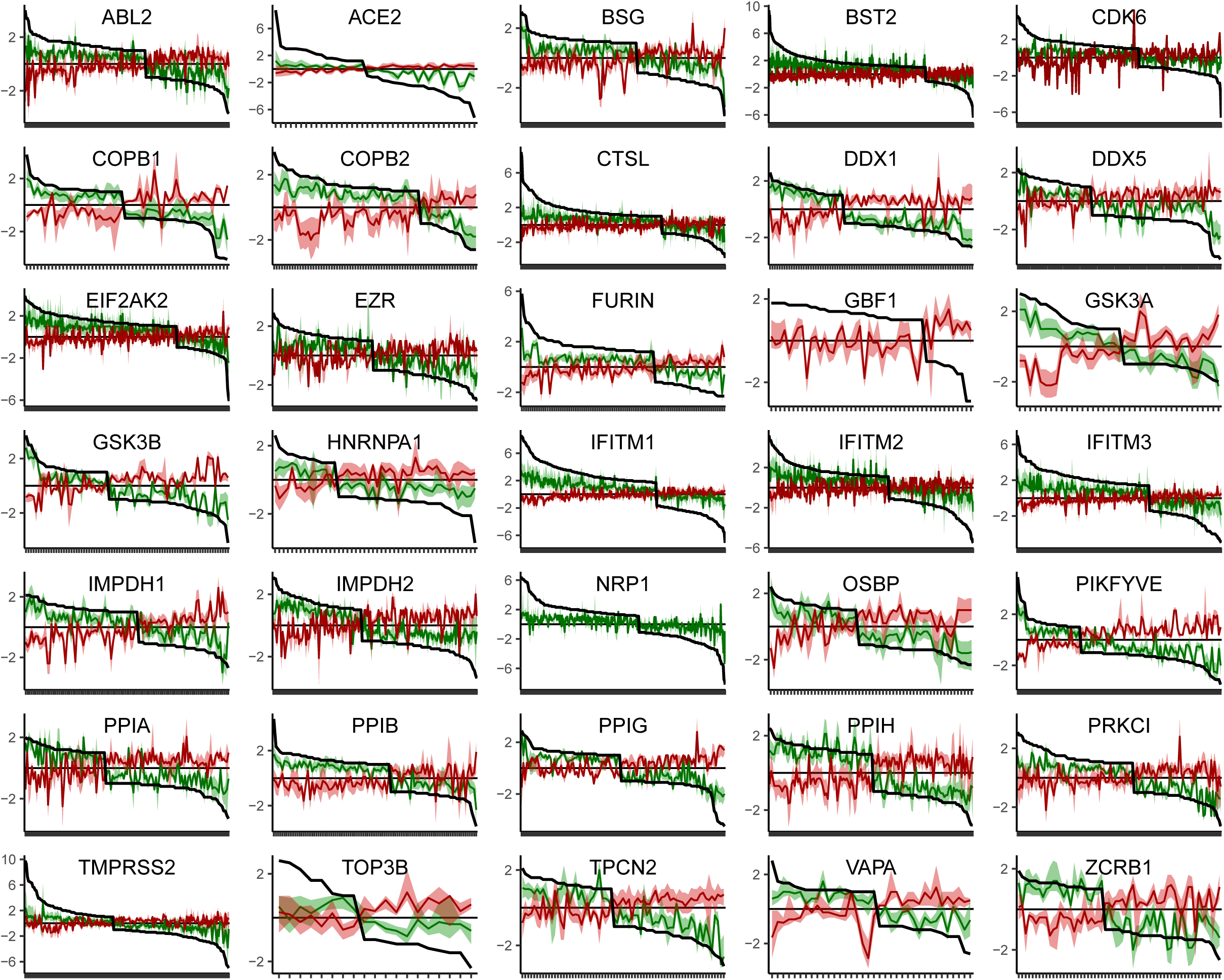
Expression profile of all seeds together with their positive and negative correlated genes. The black colored line represents the expression profile for the target gene through the different experiments (X axis), which are ordered from its higher to lower log2FC. Correlated genes are labeled in green (co-expressed genes), or red (inversely expressed genes), together with its deviation (Q1 an Q3 quartiles). Note that the number of ticks on the X axis is relative to the number of experiments for that seed.

Co-expressed genes present common functions again related to those of the seeds (Fig. 5). Pathways related of cytokine response such as interleukin or interferon signaling appear enriched to the expected antiviral genes *IFITM1, IFITM2, IFITM3, EIF2AK2*, and *BST2*, but also to the entry gene *CTSL*, and the replication genes *PPIB* and *DDX1*. In fact, the protease *CTSL* presented several interleukins as co-expressed genes, such as Interleukin-1 beta (*IL1B*), and the X-C-C motif chemokines 2 and 3 (*CXCL2, CXCL3*). Other shared pathways are those related to the cell cycle. Five genes show enrichment in cell cycle checkpoints (*DDX1, PPIB, PPIH, PRKCI*, and *ZCRB1*). The cell cycle disarrangement is an action that many viruses perform in the infected cells, where they induce a cell cycle arrest. For example, the coronavirus avian infectious bronchitis virus (IBV) activates the cell ATR signaling, which contributes to S-phase arrest and is required for efficient virus replication and progeny production (Xu et al., 2011). Genes of vesicle fusion such as *COPB1, COPB2, OSBP*, together with *PPIB*, are enriched in the expected processes of trafficking between the endoplasmic reticulum and the Golgi apparatus, glycosylation or the unfolded protein response. Other highlighted pathway is related to the nuclear export protein (NEP/NS2) of Influenza A virus that helps the transport of viral ribonucleoprotein complexes from the nucleus (Boulo et al., 2007), and it is enriched from the correlators of *DDX1, PPIB*, and *PPIH*. Other unexpected pathways were those relating *DDX1, IMPDH2, PPIH*, and *ZCRB1* to mitochondrial translation. Noteworthy, this latter pathway appeared previously related to *PPIB*, as well as the entry genes *BSG* and *TPCN2*, in the previous analysis of pathway average correlations (Fig. 3b). Finally, the most remarkable common pathway in the inversely expressed genes was interferon-alpha/beta pathway, which appeared for both *HNRNPA1* and *PPIH* genes. These genes are involved in mRNA splicing, that is one of the essential host functions for the virus, so it is expected that the cell response try to silence them, and because of this we expect that their negative correlators were related to cytokine response.

**Fig. 5.**
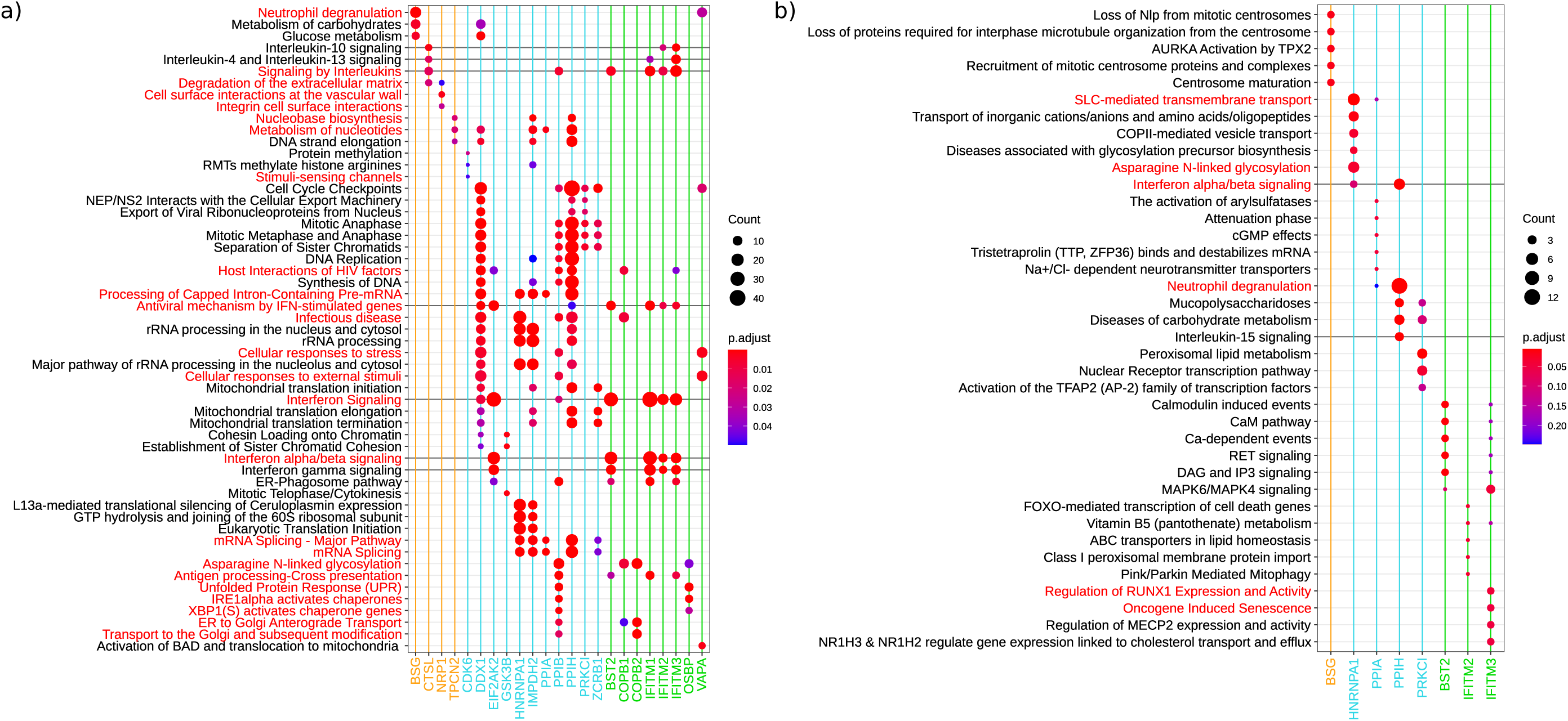
Functional enrichment of the positively (a) and negatively (b) correlated genes for all the seeds. Reactome pathways were used, with those already annotated for a seed highlighted in red color. Pathways related with interferon or interleukin signaling are highlighted with a darker line. The color of the seed name is related to their viral function (entry=orange, replication=blue, vesicle=green). Seeds without any enriched pathway are not shown.

### The best seeds’ correlators overlap with results of high-throughput experiments on SARS-CoV-2

To evaluate the list of co-expressed genes obtained, we compared them against genes identified in high-throughput laboratory experiments performed upon viral infection or in the presence of viral proteins. A recent work has completed the interactome of viral versus human proteins, and they found 332 human proteins interacting with viral proteins which are candidates to be involved in the viral infectious cycle (Gordon et al., 2020). In addition, other experimental group has published both the translatome and proteome in human cells infected by the virus (Bojkova et al., 2020), and they describe proteins that differentially change during the infection.

Two thirds of the positively correlated genes proposed by our approach appear in any of those experimental datasets (1212 out of 1899), while the expected random value would be 29 proteins in the interactome, 239 in the translatome, and 493 proteins in the proteome (Fig. 6a). However, 682 of our proposed genes did not appear as results in these experimental analyses. To assess whether these new genes could perform virus related functions their associated pathways were analyzed. In fact, a good number of genes are involved in processes related to the infectious cycle of other viruses such as Influenza A, Epstein-Barr, Hepatitis B and C, and Measles (Fig. 6b). Another function found was the homology directed repair (HDR), which is also used for the Epstein-Barr (Sugimoto et al., 2011). Furthermore, it is noteworthy the enrichment in the ubiquitin-proteasome system (Fig. 6c), since the ubiquitin mediated proteolysis is important during various stages of coronavirus infectious cycle (Raaben et al., 2010). Finally, there are functions related with the immune system which respond to virus infections such as class I MHC mediated antigen processing and presentation, and interferon/interleukin signaling.

**Fig. 6.**
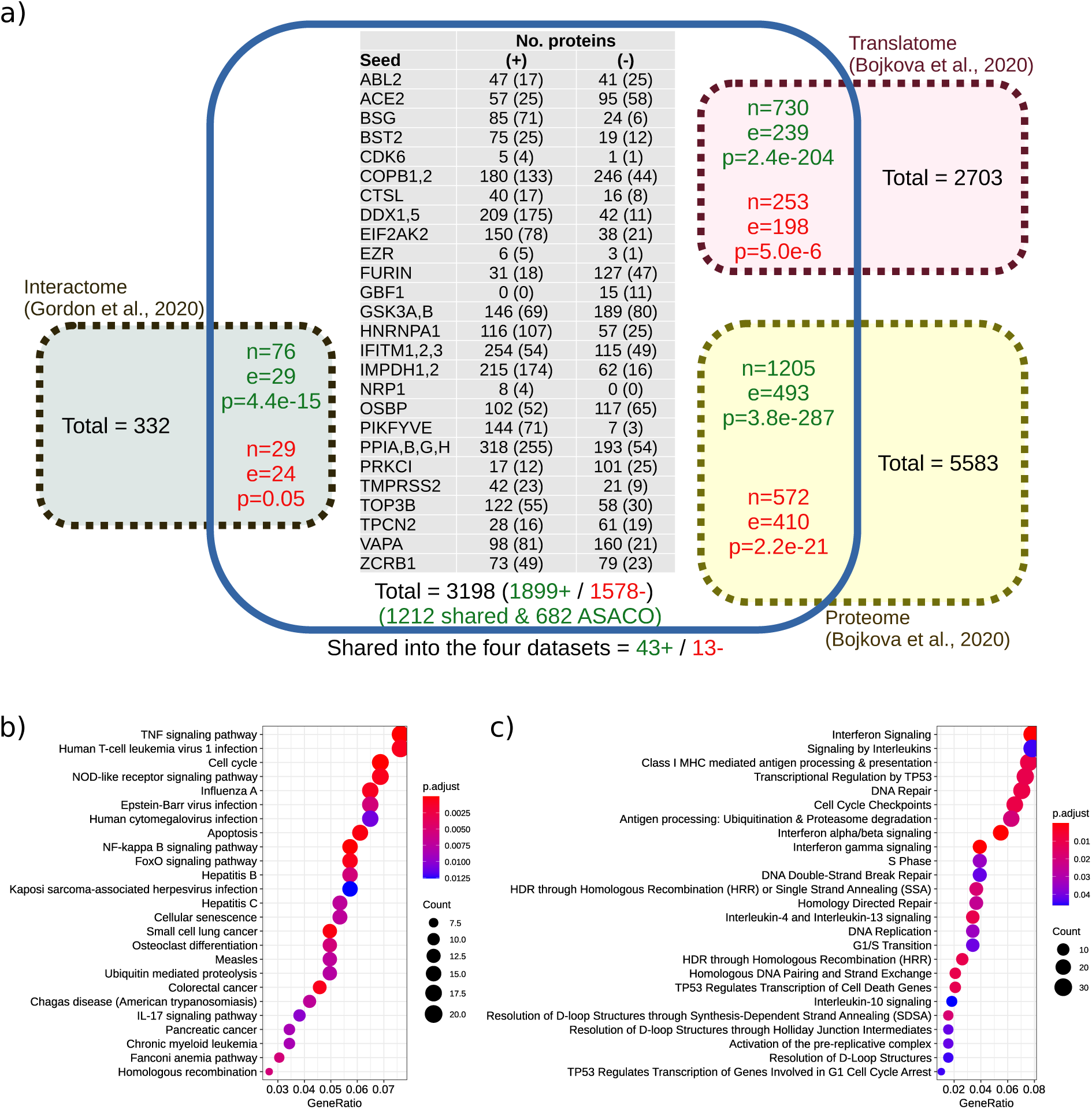
Overlapping between co-expressed proteins found by ASACO and other proteins related with the SARS-CoV-2 infection from high-throughput experiments. **a)** Co-expressed genes found by ASACO are shown inside the blue box. The interactome obtained by Gordon et al., 2020 is displayed in the green box, the translatome and the proteome by Bojkova et al., 2020 are shown in the pink and the yellow boxes respectively. The total number of proteins in each dataset is shown inside the boxes as well as the number of overlapping proteins (n), the number of coincidences expected by chance (e), and the p-value calculated with the hypergeometric distribution (p). Results for positively (green color), and negatively (red color) correlated genes are separated. The table represents the number of proteins found by ASACO for each seed, together with those that overlap with any other dataset in brackets and separated by the correlation sign. The total number of proteins is indicated below together with the total number of shared proteins and the number of positively correlated proteins exclusively found by ASACO. Outside the blue box, the number of proteins shared by the four datasets is shown. Note that the 2703 proteins in the translatome also appear in the proteome. **b)** KEGG pathway enrichment for genes exclusively found by ASACO. **c)** Reactome pathway enrichment for genes exclusively found by ASACO.

These results suggest that co-expressed genes found by our approach can offer an accurate landscape of the cellular pathways and proteins affected by the virus when the infection is progressing, even though it is based on heterogeneous experiments from many different conditions that do not include coronavirus infections.

### Transcription factors co-expressed with seeds are regulated by interferon and could induce the expression of genes involved in cell entry

The coincidence in the assigned pathways for seeds and their correlators suggests that SARS-CoV-2 host factors may belong to common regulatory networks, possibly sharing transcriptional regulators. Moreover, these regulators could be co-regulated with seeds as well. To test this hypothesis, transcription factors were identified from the correlators. So, 116 transcription factors, or putative upstream regulators, were found among the co-expressed genes, and 155 in the inversely expressed genes. Among them, 23 co-expressed regulators were common to two or more seeds (Fig. 7a). They form an interrelated network with the main antiviral genes (*EIF2AK2, BST2, IFITM1, IFITM2*, and *IFITM3*), but some of them also appeared co-expressed with proviral genes involved in vesicle fusion such as *COPB1, COPB2*, and *VAPA*, replication, as *DDX5*, and entry, suggesting common regulatory features. Related to viral entry, the transcription factor NUPR1, a stress-response protein induced by the Hepatitis B virus (Bak et al., 2015), is co-expressed with the protease TMPRSS2. Moreover, the factor ZNF267, which is an antiviral zinc finger protein (Huntley et al., 2006), is co-expressed with the kinase ABL2. Finally, TRIM14, which is a member of a family of E3 ubiquitin ligases linked to the mitochondria that plays an important role in innate defense against viruses facilitating the interferon response (Tan et al., 2017), is co-expressed with both the receptor ACE2 and the lysosomal channel TPCN2. Remarkably, all the regulators connecting the antiviral seeds with several proviral activities, including the cell entry, are genes induced by interferon. However, other transcription factors mainly connecting replication genes are contrarily repressed by interferon. For example, *YEATS4* is co-expressed with *DDX1, IMPDH2*, and *ZCRB1*, that were co-expressed with genes involved in mitochondrial translation. Other remarkable co-expressed gene is *CEBPZ*, which belongs to a family of CCAAT/enhancer-binding proteins, some of them related to immune and inflammatory response (Kinoshita et al., 1992). *CEBPZ* is a gene co-expressed with the replication seeds *DDX1* and *PPIG*, but also inversely expressed with the protease FURIN. Other correlators from this protein family are *CEBPD*, that is a co-expressed gene of IFITM3, but inversely expressed with COPB2, and *CEBPB* that is a co-expressed gene of the protease *CTSL*.

**Fig. 7.**
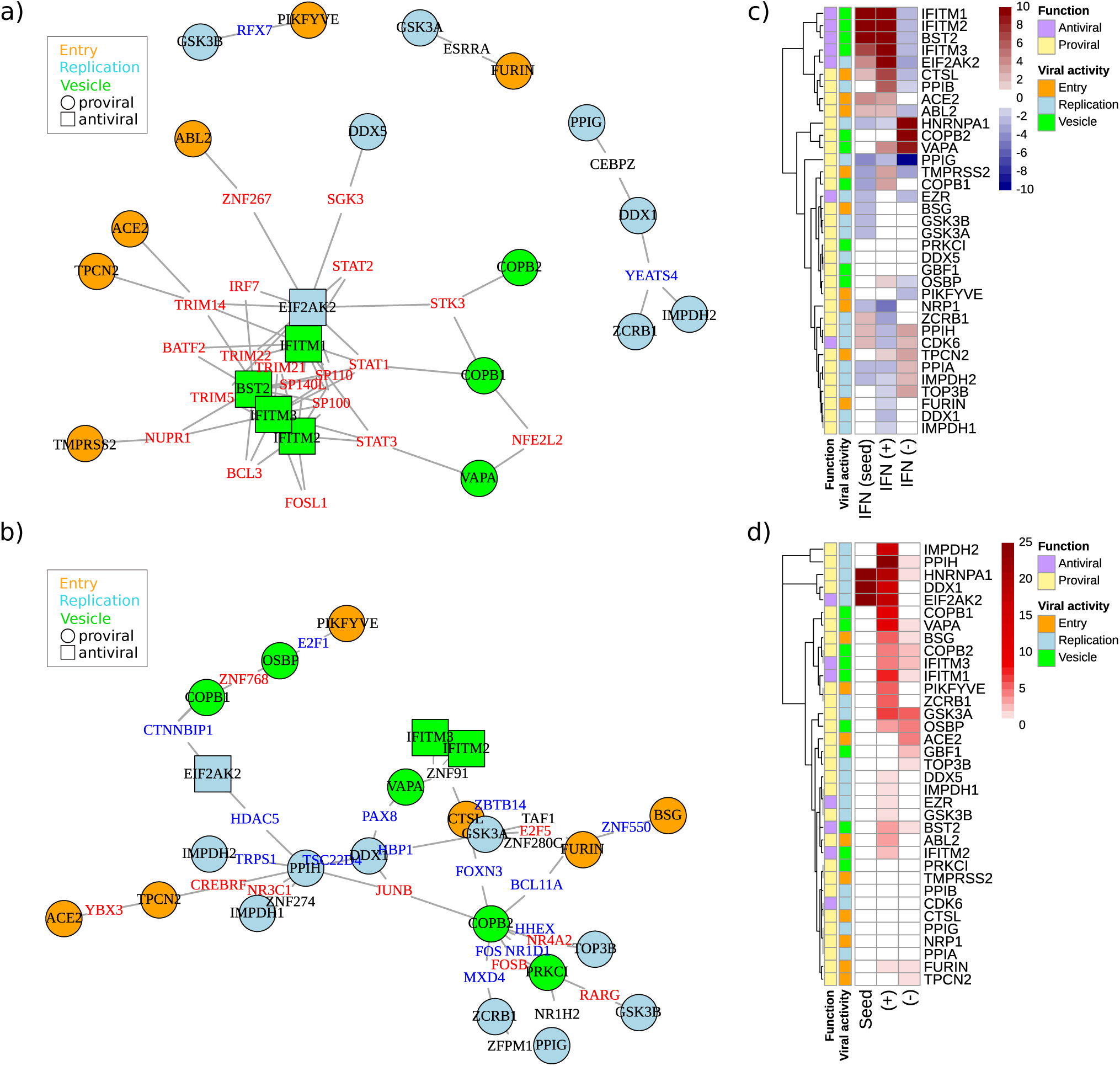
Transcription factor network from co-expressed genes, and relation with interferon and stress granules. **a)** Positively expressed transcription factors common to, at least, two seeds. Seeds are highlighted by its type of activity (proviral or antiviral), and its function in the infection (entry, replication, or vesicle fusion). Transcription factor induced by interferon are highlighted in red color and those repressed by interferon in blue color. **b)** The same as a) but for the negatively expressed transcription factors. **c)** Average interferon induced fold change of seeds, together with the value for both positively and negatively expressed genes. **d)** Number of genes related to stress granules in both positively and negatively expressed genes for each seed. The first column of the heatmap shows if the seed is related or not to stress granules.

Conversely, regulators inversely expressed with seeds are mainly interferon repressed genes (Fig. 7b). Among these genes, three zinc-finger proteins that act as antivirals against Herpex simplex virus 1 stand out in this dataset (Melchjorsen et al., 2009): *ZNF91* inversely expressed with *CTSL, IFITM2, IFITM3*, and *VAPA*, that is not induced or repressed by interferon, is a transcription factor specifically required to repress SINE-VNTR-Alu (SVA) retrotransposons (Jacobs et al., 2014); the transcription factor *ZNF550*, which is repressed by interferon and appeared inversely correlated with the entry genes *FURIN* and *BSG*; and *ZNF768*, that appeared as inversely expressed to the genes involved in vesicle fusion *COPB1* and *OSBP*. Two others inversely expressed genes were the transcriptional repressors *YBX3* and *CREBRF*, that are repressed by interferon and link the entry genes *ACE2* and *TPCN2*. Y-Box-Binding Protein 3 (YBX3) restricts Influenza A virus by impairing viral ribonucleoprotein complexes (Qin et al., 2020), and also controls amino acid levels by regulating solute carrier amino acid transporter (SLC) mRNA abundance, a pathway that is related with inversely expressed genes of *HPRNPA1* (Fig. 5b).

To further analyze the interferon signaling related with the seeds, the response to interferon was independently evaluated for each seed, together with their co-expressed and inversely expressed genes. As expected, the main interferon induced genes, the antivirals *IFITM1-3, BST2*, and *EIF2AK2*, presented an important activation together with its co-expressed genes, and a repression of their inversely expressed genes (Fig 7c). Unexpectedly, proviral genes involved in entry, *ACE2, ABL2*, and *CTSL*, together with the replication gene *PPIB*, present a similar interferon response, suggesting an undesired effect of interferon promoting SARS-CoV-2 infection. On the other hand, the seeds *HNRNPA1, COPB2*, and *VAPA* present a reversed profile. Specifically, HNRNPA1 is a gene involved in mRNA splicing, which is moved towards the stress granules during viral infection. This structures include ribonucleprotein complexes together with the cell translation machinery, and is proposedly used by the virus to perform its replication (Perdikari et al., 2020). These cytosolic particles seem to be targeted by the viral nucleocapsid protein (N). The N protein interact with 15 human proteins (Gordon et al., 2020), and 3 of them were found as co-expressed genes of *COPB1* and *COPB2* (*FAM98A*), *EIF2AK2* (*MOV10*), and *HNRNPA1* and *IMPDH2* (*PABPC4*). These three proteins have been associated to the stress granules, together with *DDX1, EIF2AK2*, and *HNRNPA1*, which reinforce the relation of the N viral protein with this stress structures (Burgess et al., 2011; Goodier et al., 2015; Ozeki et al., 2019). In fact, the antiviral seed *EIF2AK2* has been seen as the kinase activated by double-stranded RNA of viruses that activate the formation of the stress granules (Burgess and Mohr, 2018). Currently, 464 proteins are known to form part of this liquid-liquid structures, and the seed co-expressed genes include 131 of them (41 were expected by chance; 4.6e-35) (Fig. 7d), 21 from the interactome (7 expected; 7.7e-06), and 372 from the proteome (119 expected; 3.1e-137), which support the importance of these structures in the SARS-CoV-2, as well as the fact that the virus could use them for its replication and mRNA translation.

## Discussion

Currently, databases sharing gene expression data are exponentially growing due to the universalization of transcriptomics techniques (Athar et al., 2019). A secondary analysis of this data allows to reconstruct gene networks based on co-expressed genes using the so-called reverse engineering (Basso et al., 2005; He and Tan, 2016). We have developed a new *in silico* method called ASACO based on standardized gene expression data analysis. Starting from an initial seed gene, it finds the closest neighbors in terms of transcriptional regulation. Compared to other previous procedures, it has the advantage of evaluating thousands of experiments whose outcomes are normalized, allowing co-expression analysis for different heterogeneous genes over hundreds of experimental conditions (Cardozo et al., 2019; Gibbs et al., 2013). We have used this strategy on a selection of 35 cellular genes reportedly involved in SARS-CoV and/or SARS-CoV-2 infection, identifying their closest co-expressed genes. Even though the experiments employed to find them are not focused on coronaviruses infection, the functional enrichment of the co-expressed genes identify many of the already known pathways for these seeds (Fig. 3). Furthermore, both seeds and correlators fairly match most of the cellular pathways relevant to the infection cycle (Fig. 5). Moreover, these co-regulated genes show a high coincidence with the ones identified in recent high-throughput studies on cell responses to SARS-CoV-2 infection (Fig. 6a). This match can be interpreted as a cross-validation of both, experimental and *in silico* approaches, providing relevance to our identified functions and supporting the idea that co-regulation, as we identify it, can be a hallmark of co-function.

When viruses enter the host, one of the first systems that respond to the infection is the interferon signaling pathway, that induces the expression of interferon stimulated genes (ISG). Five of the seeds used in this study, considered as antiviral genes, are ISGs. *EIF2AK2* inhibits viral replication via the integrated stress response, and blocks the cellular and viral translation through the phosphorylation of EIF2α (Kang et al., 2009). *BST2* blocks the release of viruses by directly tethering nascent virions to the membranes of infected cells (Dafa-Berger et al., 2012). Finally, the transmembrane proteins *IFITM1-3* inhibit the entry of enveloped virus by preventing vesicle fusion, though they also could facilitates the infection of other viruses (Lim et al., 2016). As expected, genes positively correlated with these antiviral genes consistently belong to interferon and interleukins response pathways (Fig. 5a), and our co-expression analysis links them to regulators as STAT1, STAT3 or TRIM21 (Fig. 7ac). Surprisingly, several proviral seeds, remarkably those involved in virus entry (*ACE2, TPCN2, ABL2*, and *TMPRSS2*), are found to be linked to these antiviral genes by means of common co-expressed transcriptional regulators. The main viral receptor, ACE2, has already been found as induced by interferon, and this fact has been interpreted as “evidence that coronaviruses, as well as other viruses, have evolved to leverage features of the human IFN pathway” (Ziegler et al., 2020). This could be true for other entry genes. TRIM14 is one of the regulators found in common to *ACE2, TPCN2*, and four out of five antivirals (excluding *IFITM2*). This regulator is known to interact with MAVS at the outer mitochondria membrane and attenuates the antiviral response by the type I interferon response (Zhou et al., 2014). Since TRIM14 is co-expressed with *ACE2*, it could be responsible of, or related to, the receptor’s positive response to interferon. Interestingly, not only *ACE2* seems to be induced by interferon, but also its positive correlators, as well as those of *ABL2* (which could be induced by the ISG *ZNF267* according to our results), and the cathepsin L protease, *CTSL*. All of these are genes involved in the viral entry, and the latter activates the membrane fusion function of the spike viral protein S of SARS-CoV (Bosch et al., 2008), and could substitute to the *TMPRSS2* protease in cell types different to lung cells (Liu et al., 2020). CTSL co-expressed genes are not only interferon but also interleukin induced genes, and three cytokines are co-expressed with it (*IL1B, CXCL2*, and *CXCL3*). In addition, it shares with IFITM2, IFITM3 and the vesicle gene VAPA links to the transcriptional repressor *ZNF91*. All of this points to both ZNF91 and TRIM14 as putative regulators responsible for the presumptive interferon and interleukin dependent induction of entry proviral genes and pinpoints them as useful pharmacological targets to interfere with the viral infection. In fact, the cellular response to SARS-CoV-2 have been shown to be lightly induced by interferon, but strongly by chemokines (Blanco-Melo et al., 2020). Furthermore, CTSL appeared also linked to the co-expressed transcription factor CBPB that regulates the expression of genes involved in immune and inflammatory responses (Kinoshita et al., 1992). This transcription factor can form heterodimers with CEBPD, which is also a co-expressed gene of IFITM3, but inversely expressed gene of the vesicle protein COPB2. It suggests that the interleukin response could repress proviral genes such as *COPB2*, but undesirably induce *CTSL*.

Contrarily, genes required for viral replication such as *PPIG, DDX1, ZCRB1, IMPDH2, GSK3A*, and *GSK3B*, are bound to interferon repressed transcriptional regulators such as *RFX7* and *YEATS4*, which could negatively affect the infection in the presence of interferon.

One essential cellular pathway needed by the virus is the mRNA splicing. In this function, *HNRNPA1* is a key gene (Luo et al., 2005). Interferon triggers EIF2AK2-dependent HNRNPA1 protein translocation to stress granules resulting in mRNA translation inhibition. Until now, 464 different human proteins have been identified to participate in this liquid-liquid cytosolic structures (Nunes et al., 2019). Remarkably, proteins from stress granules show a broad convergence with our host factors co-expressed genes (131 out of 464), as well as those from the high-throughput experiments. Thus, 46 proteins out of 332 from the viral interactome are also components of stress granules. From these 46 proteins, 9 interact with the N nucleocapsid protein that seems the viral protein most related to these structures (Savastano et al., 2020). Furthermore, 20 of these proteins interact with the viral polymerase complex (nsp12, nsp7, and nsp8), and with the helicase nsp13 that in turn seems to interact with our seed *DDX5* in SARS-CoV (Chen et al., 2009a). Since virus infection leads to the hijacking of the translation machinery into the stress granules, this is a beneficial place for the translation of viral mRNA molecules, where viruses such as the respiratory syncytial virus take advantage of this (Lindquist et al., 2010). Thus, the general emergence of interferon and cytokine related genes (Blanco-Melo et al., 2020; Ziegler et al., 2020), as well as stress granules related genes (Perdikari et al., 2020; Savastano et al., 2020), revealed by this and other studies strongly suggests that SARS-CoV-2 could also use the interferon-induced stress granules as replication factory, which points to this structure as a new target for the development of therapeutic approaches to treat COVID-19.

## Conclusions

We here presented ASACO, an algorithm with the capacity to generate key functional knowledge on specific genes based on their co-expressed genes. We have tested this algorithm with host factors involved in the SARS-CoV-2 infection, by means of the analysis of co-expression data extracted from public transcriptomics databases. Although further experiments using *in vitro* and *in vivo* approaches will be required to further confirm the results obtained here, our results have allowed the discovery of relevant gene networks and cell pathways, and pointed to a series of transcription regulators as potential targets useful in the fighting against SARS-CoV-2. The consistency of our results with those obtained by other experimental approaches represent a proof of concept of the utility of this algorithm, which could be used for the study in other pathologies where there is still a need for discovering new functional knowledge.

## Acknowledgements

We thank to GaliciAME for the project that allowed the initial development of ASACO, to Gustavo Aguilar for the development of GUPO to download the transcriptomics experiments, to C3UPO for the HPC support, and to Dr. Javier Sánchez-Céspedes for the critical reading of our manuscript.

## Data and materials availability

Scripts are available in the GitHub repository, https://github.com/upobioinfo/asaco

## Author Contributions

A.J.P., A.G. and M.J.M. conceived the study, and carried out the design and coordination. A.J.P. created the *in silico* protocols and scripts and performed the analyses. G.A. made improvements to the algorithm and performed the computational analysis of the results. A.M.B., G.B., A.G. and M.J.M. analyzed and completed the results. R.R. provided the statistical support. A.J.P. and A.G. wrote the manuscript. All authors read and approved the final manuscript.

## Supplementary files

**Suppl. Fig. S1**. Common co-expressed genes among seeds.

**Suppl. Table S1**. Positively and negatively expressed genes by seed, together with their functional annotation.

## References

Amini-Bavil-Olyaee S, Choi YJ, Lee JH, Shi M, Huang IC, Farzan M, Jung JU. 2013. The antiviral effector IFITM3 disrupts intracellular cholesterol homeostasis to block viral entry. Cell Host Microbe 13:452–464. doi:10.1016/j.chom.2013.03.006

Athar A, Füllgrabe A, George N, Iqbal H, Huerta L, Ali A, Snow C, Fonseca NA, Petryszak R, Papatheodorou I, Sarkans U, Brazma A. 2019. ArrayExpress update - From bulk to single-cell expression data. Nucleic Acids Res 47:D711–D715. doi:10.1093/nar/gky964

Bak Y, Shin HJ, Bak IS, Yoon DY, Yu DY. 2015. Hepatitis B virus X promotes hepatocellular carcinoma development via nuclear protein 1 pathway. Biochem Biophys Res Commun 466:676–681. doi:10.1016/j.bbrc.2015.09.082

Basso K, Margolin AA, Stolovitzky G, Klein U, Dalla-Favera R, Califano A. 2005. Reverse engineering of regulatory networks in human B cells. Nat Genet 37:382–390. doi:10.1038/ng1532

Belouzard S, Millet JK, Licitra BN, Whittaker GR. 2012. Mechanisms of Coronavirus Cell Entry Mediated by the Viral Spike Protein. Viruses 4:1011–1033. doi:10.3390/v4061011

Blanco-Melo D, Nilsson-Payant BE, Liu WC, Uhl S, Hoagland D, Møller R, Jordan TX, Oishi K, Panis M, Sachs D, Wang TT, Schwartz RE, Lim JK, Albrecht RA, tenOever BR. 2020. Imbalanced Host Response to SARS-CoV-2 Drives Development of COVID-19. Cell 181. doi:10.1016/j.cell.2020.04.026

Bojkova D, Klann K, Koch B, Widera M, Krause D, Ciesek S, Cinatl J, Münch C. 2020. Proteomics of SARS-CoV-2-infected host cells reveals therapy targets. Nature. doi:10.1038/s41586-020-2332-7

Bosch BJ, Bartelink W, Rottier PJM. 2008. Cathepsin L Functionally Cleaves the Severe Acute Respiratory Syndrome Coronavirus Class I Fusion Protein Upstream of Rather than Adjacent to the Fusion Peptide. J Virol 82:8887–8890. doi:10.1128/jvi.00415-08

Boulant S, Stanifer M, Lozach PY. 2015. Dynamics of virus-receptor interactions in virus binding, signaling, and endocytosis. Viruses. doi:10.3390/v7062747

Boulo S, Akarsu H, Ruigrok RWH, Baudin F. 2007. Nuclear traffic of influenza virus proteins and ribonucleoprotein complexes. Virus Res. doi:10.1016/j.virusres.2006.09.013

Burgess HM, Mohr I. 2018. Defining the Role of Stress Granules in Innate Immune Suppression by the Herpes Simplex Virus 1 Endoribonuclease VHS. J Virol 92. doi:10.1128/jvi.00829-18

Burgess HM, Richardson WA, Anderson RC, Salaun C, Graham S V., Gray NK. 2011. Nuclear relocalisation of cytoplasmic poly(A)-binding proteins PABP1 and PABP4 in response to UV irradiation reveals mRNA-dependent export of metazoan PABPS. J Cell Sci 124:3344–3355. doi:10.1242/jcs.087692

Cardozo LE, Russo PST, Gomes-Correia B, Araujo-Pereira M, Sepúlveda-Hermosilla G, Maracaja-Coutinho V, Nakaya HI. 2019. WebCEMiTool: Co-expression modular analysis made easy. Front Genet 10. doi:10.3389/fgene.2019.00146

Chen JY, Chen WN, Poon KMV, Zheng BJ, Lin X, Wang YX, Wen YM. 2009a. Interaction between SARS-CoV helicase and a multifunctional cellular protein (Ddx5) revealed by yeast and mammalian cell two-hybrid systems. Arch Virol 154:507–512. doi:10.1007/s00705-009-0323-y

Chen JY, Chen WN, Poon KMV, Zheng BJ, Lin X, Wang YX, Wen YM. 2009b. Interaction between SARS-CoV helicase and a multifunctional cellular protein (Ddx5) revealed by yeast and mammalian cell two-hybrid systems. Arch Virol 154:507–512. doi:10.1007/s00705-009-0323-y

Coleman CM, Sisk JM, Mingo RM, Nelson EA, White JM, Frieman MB. 2016. Abelson Kinase Inhibitors Are Potent Inhibitors of Severe Acute Respiratory Syndrome Coronavirus and Middle East Respiratory Syndrome Coronavirus Fusion. J Virol 90:8924–8933. doi:10.1128/jvi.01429-16

Coutard B, Valle C, de Lamballerie X, Canard B, Seidah NG, Decroly E. 2020. The spike glycoprotein of the new coronavirus 2019-nCoV contains a furin-like cleavage site absent in CoV of the same clade. Antiviral Res 176:104742. doi:10.1016/j.antiviral.2020.104742

Dafa-Berger A, Kuzmina A, Fassler M, Yitzhak-Asraf H, Shemer-Avni Y, Taube R. 2012. Modulation of hepatitis C virus release by the interferon-induced protein BST-2/tetherin. Virology 428:98–111. doi:10.1016/j.virol.2012.03.011

Daly JL, Simonetti B, Antón-Plágaro C, Kavanagh Williamson M, Shoemark DK, Simón-Gracia L, Klein K, Bauer M, Hollandi R, Greber UF, Horvath P, Sessions RB, Helenius A, Hiscox JA, Teesalu T, Matthews DA, Davidson AD, Cullen PJ, Yamauchi Y. 2020. Neuropilin-1 is a host factor for SARS-CoV-2 infection. bioRxiv 2020.06.05.134114. doi:10.1101/2020.06.05.134114

de Wilde AH, Wannee KF, Scholte FEM, Goeman JJ, ten Dijke P, Snijder EJ, Kikkert M, van Hemert MJ. 2015. A Kinome-Wide Small Interfering RNA Screen Identifies Proviral and Antiviral Host Factors in Severe Acute Respiratory Syndrome Coronavirus Replication, Including Double-Stranded RNA-Activated Protein Kinase and Early Secretory Pathway Proteins. J Virol 89:8318–8333. doi:10.1128/jvi.01029-15

Durinck S, Spellman PT, Birney E, Huber W. 2009. Mapping identifiers for the integration of genomic datasets with the R/ Bioconductor package biomaRt. Nat Protoc 4:1184–1191. doi:10.1038/nprot.2009.97

Fukushi M, Yoshinaka Y, Matsuoka Y, Hatakeyama S, Ishizaka Y, Kirikae T, Sasazuki T, Miyoshi-Akiyama T. 2012. Monitoring of S Protein Maturation in the Endoplasmic Reticulum by Calnexin Is Important for the Infectivity of Severe Acute Respiratory Syndrome Coronavirus. J Virol 86:11745–11753. doi:10.1128/jvi.01250-12

Fung T, Liao Y, Liu D. 2016. Regulation of Stress Responses and Translational Control by Coronavirus. Viruses 8:184. doi:10.3390/v8070184

Fung TS, Liu DX. 2019. Human Coronavirus: Host-Pathogen Interaction. Annu Rev Microbiol 73:529–557. doi:10.1146/annurev-micro-020518-115759

Fung TS, Liu DX. 2018. Post-translational modifications of coronavirus proteins: roles and function. Future Virol 13:405–430. doi:10.2217/fvl-2018-0008

Giamarellos-Bourboulis EJ, Netea MG, Rovina N, Akinosoglou K, Antoniadou A, Antonakos N, Damoraki G, Gkavogianni T, Adami ME, Katsaounou P, Ntaganou M, Kyriakopoulou M, Dimopoulos G, Koutsodimitropoulos I, Velissaris D, Koufargyris P, Karageorgos A, Katrini K, Lekakis V, Lupse M, Kotsaki A, Renieris G, Theodoulou D, Panou V, Koukaki E, Koulouris N, Gogos C, Koutsoukou A. 2020. Complex Immune Dysregulation in COVID-19 Patients with Severe Respiratory Failure. Cell Host Microbe. doi:10.1016/j.chom.2020.04.009

Gibbs DL, Gralinski L, Baric RS, McWeeney SK. 2013. Multi-omic network signatures of disease. Front Genet 4. doi:10.3389/fgene.2013.00309

Goodier JL, Pereira GC, Cheung LE, Rose RJ, Kazazian HH. 2015. The Broad-Spectrum Antiviral Protein ZAP Restricts Human Retrotransposition. PLoS Genet 11. doi:10.1371/journal.pgen.1005252

Gordon DE, Jang GM, Bouhaddou M, Xu J, Obernier K, White KM, O’Meara MJ, Rezelj V V., Guo JZ, Swaney DL, Tummino TA, Huettenhain R, Kaake RM, Richards AL, Tutuncuoglu B, Foussard H, Batra J, Haas K, Modak M, Kim M, Haas P, Polacco BJ, Braberg H, Fabius JM, Eckhardt M, Soucheray M, Bennett MJ, Cakir M, McGregor MJ, Li Q, Meyer B, Roesch F, Vallet T, Mac Kain A, Miorin L, Moreno E, Naing ZZC, Zhou Y, Peng S, Shi Y, Zhang Z, Shen W, Kirby IT, Melnyk JE, Chorba JS, Lou K, Dai SA, Barrio-Hernandez I, Memon D, Hernandez-Armenta C, Lyu J, Mathy CJP, Perica T, Pilla KB, Ganesan SJ, Saltzberg DJ, Rakesh R, Liu X, Rosenthal SB, Calviello L, Venkataramanan S, Liboy-Lugo J, Lin Y, Huang XP, Liu YF, Wankowicz SA, Bohn M, Safari M, Ugur FS, Koh C, Savar NS, Tran QD, Shengjuler D, Fletcher SJ, O’Neal MC, Cai Y, Chang JCJ, Broadhurst DJ, Klippsten S, Sharp PP, Wenzell NA, Kuzuoglu D, Wang HY, Trenker R, Young JM, Cavero DA, Hiatt J, Roth TL, Rathore U, Subramanian A, Noack J, Hubert M, Stroud RM, Frankel AD, Rosenberg OS, Verba KA, Agard DA, Ott M, Emerman M, Jura N, von Zastrow M, Verdin E, Ashworth A, Schwartz O, d’Enfert C, Mukherjee S, Jacobson M, Malik HS, Fujimori DG, Ideker T, Craik CS, Floor SN, Fraser JS, Gross JD, Sali A, Roth BL, Ruggero D, Taunton J, Kortemme T, Beltrao P, Vignuzzi M, García-Sastre A, Shokat KM, Shoichet BK, Krogan NJ. 2020. A SARS-CoV-2 protein interaction map reveals targets for drug repurposing. Nature. doi:10.1038/s41586-020-2286-9

He B, Tan K. 2016. Understanding transcriptional regulatory networks using computational models, Current Opinion in Genetics and Development. Elsevier Ltd. doi:10.1016/j.gde.2016.02.002

Heald-Sargent T, Gallagher T. 2012. Ready, Set, Fuse! The Coronavirus Spike Protein and Acquisition of Fusion Competence. Viruses 4:557–580. doi:10.3390/v4040557

Hoffmann M, Kleine-Weber H, Schroeder S, Mü MA, Drosten C, Pö S, Krü N, Herrler T, Erichsen S, Schiergens TS, Herrler G, Wu N-H, Nitsche A, Pö Hlmann S. 2020. SARS-CoV-2 Cell Entry Depends on ACE2 and TMPRSS2 and Is Blocked by a Clinically Proven Protease Inhibitor Article SARS-CoV-2 Cell Entry Depends on ACE2 and TMPRSS2 and Is Blocked by a Clinically Proven Protease Inhibitor. Cell 181:1–10. doi:10.1016/j.cell.2020.02.052

Huang C, Wang Y, Li X, Ren L, Zhao J, Hu Y, Zhang L, Fan G, Xu J, Gu X, Cheng Z, Yu T, Xia J, Wei Y, Wu W, Xie X, Yin W, Li H, Liu M, Xiao Y, Gao H, Guo L, Xie J, Wang G, Jiang R, Gao Z, Jin Q, Wang J, Cao B. 2020. Clinical features of patients infected with 2019 novel coronavirus in Wuhan, China. Lancet 395:497–506. doi:10.1016/S0140-6736(20)30183-5

Huang I-C, Bailey CC, Weyer JL, Radoshitzky SR, Becker MM, Chiang JJ, Brass AL, Ahmed AA, Chi X, Dong L, Longobardi LE, Boltz D, Kuhn JH, Elledge SJ, Bavari S, Denison MR, Choe H, Farzan M. 2011. Distinct Patterns of IFITM-Mediated Restriction of Filoviruses, SARS Coronavirus, and Influenza A Virus. PLoS Pathog 7:e1001258. doi:10.1371/journal.ppat.1001258

Huntley S, Baggott DM, Hamilton AT, Tran-Gyamfi M, Yang S, Kim J, Gordon L, Branscomb E, Stubbs L. 2006. A comprehensive catalog of human KRAB-associated zinc finger genes: Insights into the evolutionary history of a large family of transcriptional repressors. Genome Res 16:669–677. doi:10.1101/gr.4842106

Jacobs FMJ, Greenberg D, Nguyen N, Haeussler M, Ewing AD, Katzman S, Paten B, Salama SR, Haussler D. 2014. An evolutionary arms race between KRAB zinc-finger genes ZNF91/93 and SVA/L1 retrotransposons. Nature 516:242–245. doi:10.1038/nature13760

Jassal B, Matthews L, Viteri G, Gong C, Lorente P, Fabregat A, Sidiropoulos K, Cook J, Gillespie M, Haw R, Loney F, May B, Milacic M, Rothfels K, Sevilla C, Shamovsky V, Shorser S, Varusai T, Weiser J, Wu G, Stein L, Hermjakob H, D’Eustachio P. 2020. The reactome pathway knowledgebase. Nucleic Acids Res 48:D498–D503. doi:10.1093/nar/gkz1031

Kamitani W, Huang C, Narayanan K, Lokugamage KG, Makino S. 2009. A two-pronged strategy to suppress host protein synthesis by SARS coronavirus Nsp1 protein. Nat Struct Mol Biol 16:1134–1140. doi:10.1038/nsmb.1680

Kanehisa M, Furumichi M, Tanabe M, Sato Y, Morishima K. 2017. KEGG: new perspectives on genomes, pathways, diseases and drugs. Nucleic Acids Res 45:D353–D361. doi:10.1093/nar/gkw1092

Kang J Il, Kwon SN, Park SH, Kim YK, Choi SY, Kim JP, Ahn BY. 2009. PKR protein kinase is activated by hepatitis C virus and inhibits viral replication through translational control. Virus Res 142:51–56. doi:10.1016/j.virusres.2009.01.007

Kinoshita S, Akira S, Kishimoto T. 1992. A member of the C/EBP family, NF-IL6β, forms a heterodimer and transcriptionally synergizes with NF-IL6. Proc Natl Acad Sci U S A 89:1473–1476. doi:10.1073/pnas.89.4.1473

Lim Y, Ng Y, Tam J, Liu D. 2016. Human Coronaviruses: A Review of Virus–Host Interactions. Diseases 4:26. doi:10.3390/diseases4030026

Lindquist ME, Lifland AW, Utley TJ, Santangelo PJ, Crowe JE. 2010. Respiratory Syncytial Virus Induces Host RNA Stress Granules To Facilitate Viral Replication. J Virol 84:12274–12284. doi:10.1128/jvi.00260-10

Liu H, Gai S, Wang X, Zeng J, Sun C, Zhao Y, Zheng Z. 2020. Single-cell analysis of SARS-CoV-2 receptor ACE2 and spike protein priming expression of proteases in the human heart. Cardiovasc Res. doi:10.1093/cvr/cvaa191

Lobo I. 2008. Biological Complexity and Integrative Levels of Organization. Nat Educ.

Long HK, Prescott SL, Wysocka J. 2016. Ever-Changing Landscapes: Transcriptional Enhancers in Development and Evolution. Cell. doi:10.1016/j.cell.2016.09.018

Luo H, Chen Q, Chen J, Chen K, Shen X, Jiang H. 2005. The nucleocapsid protein of SARS coronavirus has a high binding affinity to the human cellular heterogeneous nuclear ribonucleoprotein A1. FEBS Lett 579:2623–2628. doi:10.1016/j.febslet.2005.03.080

Masters PS. 2006. The Molecular Biology of Coronaviruses. Adv Virus Res. doi:10.1016/S0065-3527(06)66005-3

Melchjorsen J, Matikainen S, Paludan SR. 2009. Activation and evasion of innate antiviral immunity by herpes simplex virus. Viruses. doi:10.3390/v1030737

Millet JK, Kien F, Cheung C-YY, Siu Y-LL, Chan W-LL, Li H, Leung H-LL, Jaume M, Bruzzone R, Malik Peiris JS, Altmeyer RM, Nal B. 2012. Ezrin Interacts with the SARS Coronavirus Spike Protein and Restrains Infection at the Entry Stage. PLoS One 7:e49566. doi:10.1371/journal.pone.0049566

Moosa MM, Banerjee PR. 2020. Subversion of host stress granules by coronaviruses: Potential roles of π-rich disordered domains of viral nucleocapsids. J Med Virol jmv.26195. doi:10.1002/jmv.26195

Nunes C, Mestre I, Marcelo A, Koppenol R, Matos CA, Nóbrega C. 2019. MSGP: The first database of the protein components of the mammalian stress granules. Database 2019. doi:10.1093/database/baz031

Ou X, Liu Y, Lei X, Li P, Mi D, Ren L, Guo L, Guo R, Chen T, Hu J, Xiang Z, Mu Z, Chen X, Chen J, Hu K, Jin Q, Wang J, Qian Z. 2020. Characterization of spike glycoprotein of SARS-CoV-2 on virus entry and its immune cross-reactivity with SARS-CoV. Nat Commun 11. doi:10.1038/s41467-020-15562-9

Ozeki K, Sugiyama M, Akter KA, Nishiwaki K, Asano-Inami E, Senga T. 2019. FAM98A is localized to stress granules and associates with multiple stress granule-localized proteins. Mol Cell Biochem 451:107–115. doi:10.1007/s11010-018-3397-6

Papatheodorou I, Moreno P, Manning J, Fuentes AM-P, George N, Fexova S, Fonseca NA, Füllgrabe A, Green M, Huang N, Huerta L, Iqbal H, Jianu M, Mohammed S, Zhao L, Jarnuczak AF, Jupp S, Marioni J, Meyer K, Petryszak R, Prada Medina CA, Talavera-López C, Teichmann S, Vizcaino JA, Brazma A. 2020. Expression Atlas update: from tissues to single cells. Nucleic Acids Res 48:D77–D83. doi:10.1093/nar/gkz947

Perdikari TM, Murthy AC, Ryan VH, Watters S, Naik MT, Fawzi NL. 2020. SARS-CoV-2 nucleocapsid protein undergoes liquid-liquid phase separation stimulated by RNA and partitions into phases of human ribonucleoproteins. bioRxiv 2020.06.09.141101. doi:10.1101/2020.06.09.141101

Pfefferle S, Schöpf J, Kögl M, Friedel CC, Müller MA, Carbajo-Lozoya J, Stellberger T, von Dall’Armi E, Herzog P, Kallies S, Niemeyer D, Ditt V, Kuri T, Züst R, Pumpor K, Hilgenfeld R, Schwarz F, Zimmer R, Steffen I, Weber F, Thiel V, Herrler G, Thiel H-J, Schwegmann-Weßels C, Pöhlmann S, Haas J, Drosten C, von Brunn A. 2011. The SARS-Coronavirus-Host Interactome: Identification of Cyclophilins as Target for Pan-Coronavirus Inhibitors. PLoS Pathog 7:e1002331. doi:10.1371/journal.ppat.1002331

Pinto BG, Oliveira AE, Singh Y, Jimenez L, Goncalves AN, Ogava RL, Creighton R, Peron JP, Nakaya HI. 2020. ACE2 Expression is Increased in the Lungs of Patients with Comorbidities Associated with Severe COVID-19. medRxiv 2020.03.21.20040261. doi:10.1101/2020.03.21.20040261

Prasanth KR, Hirano M, Fagg WS, McAnarney ET, Shan C, Xie X, Hage A, Pietzsch CA, Bukreyev A, Rajsbaum R, Shi P-Y, Bedford MT, Bradrick SS, Menachery V, Garcia-Blanco MA. 2020. Topoisomerase III-ß is required for efficient replication of positive-sense RNA viruses. bioRxiv 2020.03.24.005900. doi:10.1101/2020.03.24.005900

Qin Z, Qu X, Lei L, Xu L, Pan Z. 2020. Y-box-binding protein 3 (YBX3) restricts influenza A virus by interacting with viral ribonucleoprotein complex and imparing its function. J Gen Virol 101:385–398. doi:10.1099/jgv.0.001390

Raaben M, Posthuma CC, Verheije MH, te Lintelo EG, Kikkert M, Drijfhout JW, Snijder EJ, Rottier PJM, de Haan CAM. 2010. The Ubiquitin-Proteasome System Plays an Important Role during Various Stages of the Coronavirus Infection Cycle. J Virol 84:7869–7879. doi:10.1128/jvi.00485-10

Rogers JM, Bulyk ML. 2018. Diversification of transcription factor–DNA interactions and the evolution of gene regulatory networks. Wiley Interdiscip Rev Syst Biol Med. doi:10.1002/wsbm.1423

Rusinova I, Forster S, Yu S, Kannan A, Masse M, Cumming H, Chapman R, Hertzog PJ. 2013. INTERFEROME v2.0: An updated database of annotated interferon-regulated genes. Nucleic Acids Res 41. doi:10.1093/nar/gks1215

Saijo M, Morikawa S, Fukushi S, Mizutani T, Hasegawa H, Nagata N, Iwata N, Kurane I. 2005. Inhibitory effect of mizoribine and ribavirin on the replication of severe acute respiratory syndrome (SARS)-associated coronavirus. Antiviral Res 66:159–163. doi:10.1016/j.antiviral.2005.01.003

Sanche S, Lin YT, Xu C, Romero-Severson E, Hengartner N, Ke R. 2020. High Contagiousness and Rapid Spread of Severe Acute Respiratory Syndrome Coronavirus 2. Emerg Infect Dis 26. doi:10.3201/eid2607.200282

Savastano A, Opakua AI de, Rankovic M, Zweckstetter M. 2020. Nucleocapsid protein of SARS-CoV-2 phase separates into RNA-rich polymerase-containing condensates. bioRxiv 2020.06.18.160648. doi:10.1101/2020.06.18.160648

Sola I, Galan C, Mateos-Gomez PA, Palacio L, Zuniga S, Cruz JL, Almazan F, Enjuanes L. 2011. The Polypyrimidine Tract-Binding Protein Affects Coronavirus RNA Accumulation Levels and Relocalizes Viral RNAs to Novel Cytoplasmic Domains Different from Replication-Transcription Sites. J Virol 85:5136–5149. doi:10.1128/jvi.00195-11

Spitz F, Furlong EEM. 2012. Transcription factors: From enhancer binding to developmental control. Nat Rev Genet. doi:10.1038/nrg3207

Sugimoto A, Kanda T, Yamashita Y, Murata T, Saito S, Kawashima D, Isomura H, Nishiyama Y, Tsurumi T. 2011. Spatiotemporally different DNA repair systems participate in Epstein-Barr virus genome maturation. J Virol 85:6127–35. doi:10.1128/JVI.00258-11

Sungnak W, Huang N, Bécavin C, Berg M, Queen R, Litvinukova M, Talavera-López C, Maatz H, Reichart D, Sampaziotis F, Worlock KB, Yoshida M, Barnes JL, Banovich NE, Barbry P, Brazma A, Collin J, Desai TJ, Duong TE, Eickelberg O, Falk C, Farzan M, Glass I, Gupta RK, Haniffa M, Horvath P, Hubner N, Hung D, Kaminski N, Krasnow M, Kropski JA, Kuhnemund M, Lako M, Lee H, Leroy S, Linnarson S, Lundeberg J, Meyer KB, Miao Z, Misharin A V., Nawijn MC, Nikolic MZ, Noseda M, Ordovas-Montanes J, Oudit GY, Pe’er D, Powell J, Quake S, Rajagopal J, Tata PR, Rawlins EL, Regev A, Reyfman PA, Rozenblatt-Rosen O, Saeb-Parsy K, Samakovlis C, Schiller HB, Schultze JL, Seibold MA, Seidman CE, Seidman JG, Shalek AK, Shepherd D, Spence J, Spira A, Sun X, Teichmann SA, Theis FJ, Tsankov AM, Vallier L, van den Berge M, Whitsett J, Xavier R, Xu Y, Zaragosi LE, Zerti D, Zhang H, Zhang K, Rojas M, Figueiredo F. 2020. SARS-CoV-2 entry factors are highly expressed in nasal epithelial cells together with innate immune genes. Nat Med 26:681–687. doi:10.1038/s41591-020-0868-6

Tan P, He L, Cui J, Qian C, Cao X, Lin M, Zhu Q, Li Y, Xing C, Yu X, Wang HY, Wang RF. 2017. Assembly of the WHIP-TRIM14-PPP6C Mitochondrial Complex Promotes RIG-I-Mediated Antiviral Signaling. Mol Cell 68:293-307.e5. doi:10.1016/j.molcel.2017.09.035

Tan YW, Hong W, Liu DX. 2012. Binding of the 5 0-untranslated region of coronavirus RNA to zinc finger CCHC-type and RNA-binding motif 1 enhances viral replication and transcription. Nucleic Acids Res 40:5065–77. doi:10.1093/nar/gks165

Taylor JK, Coleman CM, Postel S, Sisk JM, Bernbaum JG, Venkataraman T, Sundberg EJ, Frieman MB. 2015. Severe Acute Respiratory Syndrome Coronavirus ORF7a Inhibits Bone Marrow Stromal Antigen 2 Virion Tethering through a Novel Mechanism of Glycosylation Interference. J Virol 89:11820–11833. doi:10.1128/jvi.02274-15

Thoms M, Buschauer R, Ameismeier M, Koepke L, Denk T, Hirschenberger M, Kratzat H, Hayn M, Mackens-Kiani T, Cheng J, Stuerzel CM, Froehlich T, Berninghausen O, Becker T, Kirchhoff F, Sparrer KMJ, Beckmann R. 2020. Structural basis for translational shutdown and immune evasion by the Nsp1 protein of SARS-CoV-2. bioRxiv 2020.05.18.102467. doi:10.1101/2020.05.18.102467

Wang K, Chen W, Zhou Y-S, Lian J-Q, Zhang Z, Du P, Gong L, Zhang Y, Cui H-Y, Geng J-J, Wang B, Sun X-X, Wang C-F, Yang X-MX, Lin P, Deng Y-Q, Wei D, Yang X-MX, Zhu Y-M, Zhang K, Zheng Z-H, Miao J-L, Guo T, Shi Y, Zhang J, Fu L, Wang Q-Y, Bian H, Zhu P, Chen Z-N. 2020. SARS-CoV-2 invades host cells via a novel route: CD147-spike protein. bioRxiv 2020.03.14.988345. doi:10.1101/2020.03.14.988345

Wu CH, Chen PJ, Yeh SH. 2014. Nucleocapsid phosphorylation and RNA helicase DDX1 recruitment enables coronavirus transition from discontinuous to continuous transcription. Cell Host Microbe 16:462–472. doi:10.1016/j.chom.2014.09.009

Wu CH, Yeh SH, Tsay YG, Shieh YH, Kao CL, Chen YS, Wang SH, Kuo TJ, Chen DS, Chen PJ. 2009. Glycogen synthase kinase-3 regulates the phosphorylation of severe acute respiratory syndrome coronavirus mucleocapsid protein and viral replication. J Biol Chem 284:5229–5239. doi:10.1074/jbc.M805747200

Xu LH, Huang M, Fang SG, Liu DX. 2011. Coronavirus infection induces DNA replication stress partly through interaction of its nonstructural protein 13 with the p125 subunit of DNA polymerase δ. J Biol Chem 286:39546–39559. doi:10.1074/jbc.M111.242206

Zavašnik-Bergant V, Schweiger A, Bevec T, Golouh R, Turk V, Kos J. 2004. Inhibitory p41 isoform of invariant chain and its potential target enzymes cathepsins L and H in distinct populations of macrophages in human lymph nodes. Immunology 112:378–385. doi:10.1111/j.1365-2567.2004.01879.x

Zhou P, Yang X Lou, Wang XG, Hu B, Zhang L, Zhang W, Si HR, Zhu Y, Li B, Huang CL, Chen HD, Chen J, Luo Y, Guo H, Jiang R Di, Liu MQ, Chen Y, Shen XR, Wang XG, Zheng XS, Zhao K, Chen QJ, Deng F, Liu LL, Yan B, Zhan FX, Wang YY, Xiao GF, Shi ZL. 2020. A pneumonia outbreak associated with a new coronavirus of probable bat origin. Nature 579:270–273. doi:10.1038/s41586-020-2012-7

Zhou Yonggang, Fu B, Zheng X, Wang D, Zhao C, Qi Y, Sun R, Tian Z, Xu X, Wei H. 2020. Pathogenic T-cells and inflammatory monocytes incite inflammatory storms in severe COVID-19 patients, Natl Sci Rev.

Zhou Yadi, Hou Y, Shen J, Huang Y, Martin W, Cheng F. 2020. Network-based drug repurposing for novel coronavirus 2019-nCoV/SARS-CoV-2. Cell Discov 6. doi:10.1038/s41421-020-0153-3

Zhou Z, Jia X, Xue Q, Dou Z, Ma Y, Zhao Z, Jiang Z, He B, Jin Q, Wang J. 2014. TRIM14 is a mitochondrial adaptor that facilitates retinoic acid-inducible gene-I-like receptor-mediatedinnate immune response. Proc Natl Acad Sci U S A 111. doi:10.1073/pnas.1316941111

Ziegler CGK, Allon SJ, Nyquist SK, Mbano IM, Miao VN, Tzouanas CN, Cao Y, Yousif AS, Bals J, Hauser BM, Feldman J, Muus C, Wadsworth MH, Kazer SW, Hughes TK, Doran B, Gatter GJ, Vukovic M, Taliaferro F, Mead BE, Guo Z, Wang JP, Gras D, Plaisant M, Ansari M, Angelidis I, Adler H, Sucre JMS, Taylor CJ, Lin B, Waghray A, Mitsialis V, Dwyer DF, Buchheit KM, Boyce JA, Barrett NA, Laidlaw TM, Carroll SL, Colonna L, Tkachev V, Peterson CW, Yu A, Zheng HB, Gideon HP, Winchell CG, Lin PL, Bingle CD, Snapper SB, Kropski JA, Theis FJ, Schiller HB, Zaragosi LE, Barbry P, Leslie A, Kiem HP, Flynn JAL, Fortune SM, Berger B, Finberg RW, Kean LS, Garber M, Schmidt AG, Lingwood D, Shalek AK, Ordovas-Montanes J, Banovich N, Brazma A, Desai T, Duong TE, Eickelberg O, Falk C, Farzan M, Glass I, Haniffa M, Horvath P, Hung D, Kaminski N, Krasnow M, Kuhnemund M, Lafyatis R, Lee H, Leroy S, Linnarson S, Lundeberg J, Meyer K, Misharin A, Nawijn M, Nikolic MZ, Pe’er D, Powell J, Quake S, Rajagopal J, Tata PR, Rawlins EL, Regev A, Reyfman PA, Rojas M, Rosen O, Saeb-Parsy K, Samakovlis C, Schiller H, Schultze JL, Seibold MA, Shepherd D, Spence J, Spira A, Sun X, Teichmann S, Theis F, Tsankov A, van den Berge M, von Papen M, Whitsett J, Xavier R, Xu Y, Zhang K. 2020. SARS-CoV-2 Receptor ACE2 Is an Interferon-Stimulated Gene in Human Airway Epithelial Cells and Is Detected in Specific Cell Subsets across Tissues. Cell 181. doi:10.1016/j.cell.2020.04.035

