## Supplementary figures and images for "Serial co-expression analysis of host factors from SARS-CoV viruses highly converges with former high-throughput screenings and proposes key regulators and co-option of cellular pathways"

### Suppl. Fig. S1

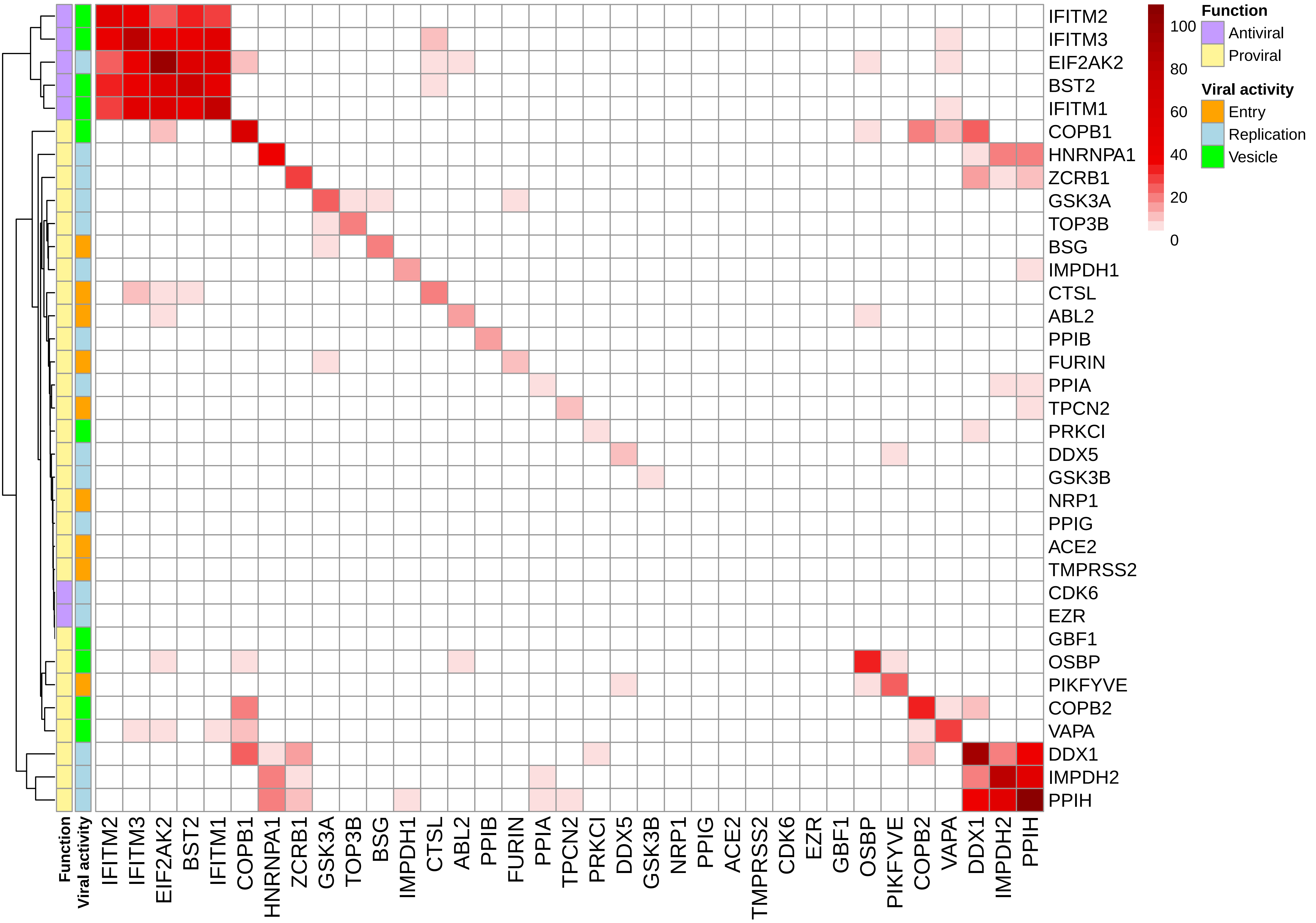
